# Bidirectionally responsive thermoreceptors encode cool and warm

**DOI:** 10.1101/2024.11.28.625856

**Authors:** Phillip Bokiniec, Clarissa J. Whitmire, James F.A. Poulet

**Affiliations:** Max Delbrück Center for Molecular Medicine in the Helmholtz Association (MDC), Berlin, Germany; Neuroscience Research Center, Charité-Universitätsmedizin Berlin, Berlin, Germany; The University of Queensland, Queensland Brain Institute, St Lucia, Brisbane, Queensland, Australia; The University of Queensland, School of Electrical Engineering and Computer Science, St Lucia, Brisbane, Queensland, Australia

## Abstract

Thermal sensation is a fundamental sense initiated by the activity of primary afferent thermoreceptors. While considerable attention has been paid to the encoding of noxious temperatures by thermoreceptors, it is far less clear how they encode innocuous cool and warm which are more commonly encountered in the environment. To address this, we sampled the entire thermoreceptor population using *in vivo* two-photon calcium imaging in the lumbar dorsal root ganglia of awake and anesthetized mice. We found that the vast majority of thermoreceptors respond bidirectionally, with an enhanced response to cool and a suppressed response to warm. Using *in vivo* pharmacology and computational modelling, we demonstrate that conductance changes in the cool-sensitive TRPM8 channel are sufficient to explain this bidirectional response type. Our comprehensive dataset reveals the fundamental principles of the peripheral encoding of innocuous temperatures and suggests that the same population of thermoreceptors underlie the distinct sensations of cool and warm.

## Introduction

The temperature of a surface provides important information about its identity and is most often experienced in an innocuous range. Detecting environmental temperature relies on the activity of specialized primary afferent neurons, or ‘thermoreceptors’ that innervate the skin, and whose cell bodies are located in the trigeminal or dorsal root ganglia (TRG, DRG). As one of the most accessible cell types in the nervous system, thermoreceptors have been examined for over a century^1–4^, yet the way in which innocuous cool and warm are encoded by thermoreceptors remains unclear.

Distinct characteristics of warm and cool, including a higher perceptual threshold for warm^5–7^ and a stronger representation of cool in the nervous system^8–15^, suggest that the afferent encoding of warm and cool is functionally and anatomically distinct. A simple model, for example, is that different thermoreceptors show an increase in firing rate to either warm or cool^16,17^. This aligns with the finding of discrete areas of skin that mediate cool or warm sensations^2^, and the identification of thermally sensitive transient receptor potential (TRP) channels that are activated within specific temperature ranges^18–21^. However, the cool sensitive TRPM8 channel has been shown to contribute to the perception of both cool and warm^7,22^, and thermoreceptors have been identified with ongoing firing that exhibit bidirectional increases and decreases in response to transient changes in temperature in humans, primates, mice, and Drosophila^7,8,16,23–26^. Overall, the fundamental question of whether thermoreceptor encoding of warm and cool is integrated or separate remains unresolved.

In part, this is because most *in vivo* efforts to address cool and warm encoding by thermoreceptors relied on electrophysiological single unit recordings^7,8,10,25,27,28^. While these studies provided valuable information, they were constrained by small sample sizes and relatively short recording durations that allowing limited probing of possible stimulus space. *In vivo* 2-photon imaging now provides more stable access to larger numbers of genetically identified populations of neurons, but prior work using this approach has focused on the encoding of thermal pain, the impact of inflammation, or thermoreceptor genetic identity^17,29–32^ and not reached a consensus on the encoding of innocuous temperatures.

Here we take advantage of a mouse line that labels all thermoreceptors (TRPV1-cre^33^), and use an *in vivo* two-photon calcium imaging preparation that allows chronic access to large numbers of lumbar DRG thermoreceptors in awake and anesthetized mice. Combining imaging with *in vivo* pharmacological manipulations and computational modelling allowed us to define how mammalian sensory afferent neurons encode innocuous temperatures.

## Results

### Thermoreceptor encoding of cool and warm in awake mice

The TRPV1 lineage of DRG neurons is required for thermosensation and provides a molecular target for all thermoreceptors^33^. Using mice expressing GCaMP6 in thermoreceptors (*TRPV1::cre;Ai162d* and *TRPV1::cre;Ai148d*), we performed a comprehensive investigation into the primary afferent encoding of innocuous temperature. To image the activity of GCaMP6 expressing thermoreceptors that innervate the hindpaw, we established a chronic glass window preparation that allowed optical access to the lumbar 4 (L4) DRG^34^ (**Fig. 1a**). Awake mice were then habituated to spinal clamp fixation and thermal stimuli were applied to the plantar surface of the hindpaw during two-photon calcium imaging.

**Figure 1.**
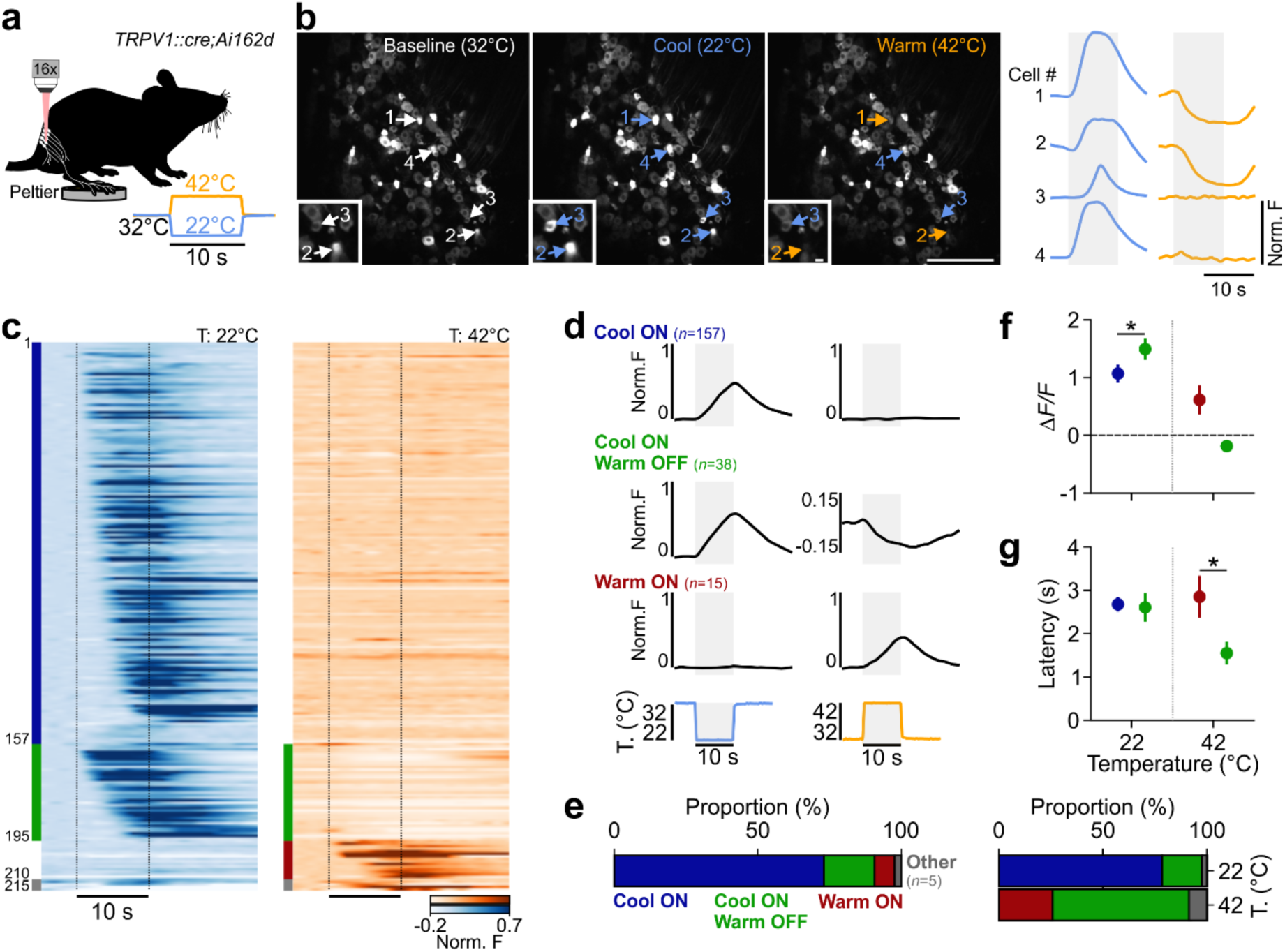
Thermoreceptor encoding of cool and warm in the awake mouse. **a.** Schematic of an awake TRPV1-GCamP6s mouse with right hindpaw tethered to a Peltier element for cutaneous thermal stimulation during two-photon calcium imaging of the right L4 DRG; inset shows temporal dynamics of cooling and warming stimuli. **b.** Left, example of *in vivo* two-photon images of L4 DRG at an adapted temperature (AT) of 32°C, during a cooling stimulus (22°C), and during a warming stimulus (42°C). Scale bar, 300 µm. Inset shows example of two thermoreceptors, scale bar, 25 µm. Right, example mean responses of single thermoreceptors during a 10°C cooling (blue) or 10°C warming (orange) stimulus from an AT of 32°C (*n =* 10 trials each). Arrows and numbers identify single neurons labelled in images. Shaded gray box indicates length of stimulus (10 s). **c.** Single thermoreceptor calcium responses to cooling (left) and warming (right) stimuli. Vertical dashed lines indicate the onset and end of the plateau phase of a thermal stimulus (*n =* 215 thermally responsive neurons from 12 awake mice). Index bar on the left indicates thermal functional response type as indicated in panel d. **d.** Grand average responses to cooling (left) or warming (right) stimuli, of thermoreceptors showing either an ON response to cooling (blue, *n =* 157), an ON response to cooling and an OFF response to warming (green, *n =* 38), or an ON response to warming (maroon, *n =* 15), shaded gray box indicates length of stimulus (10 s). **e.** Left, proportion of functional types of thermoreceptors (*n =* 157 Cool-ON, blue; *n =* 38 Cool-ON/Warm-OFF, green; *n =* 15 Warm-ON, maroon; *n =* 5 Other, grey). Right, proportion of cool (top) and warm (bottom) responses from all thermoreceptors. **f.** Peak response amplitudes for cool (left) and warm (right) responses from each response classification as indicated in panel d. **g.** Onset latencies for cool (left) and warm (right) responses from each response classification as indicated in panel d. All data values show mean ± sem.

First, we delivered Δ10°C cool and warm stimuli (10 s duration, 75°C/s ramp speed) via a Peltier element from an adapted temperature (AT) of 32°C. Throughout this study, cool stimuli are those at lower temperatures than the AT, whereas warm stimuli are those at higher values than the AT. A sparse population of cells displayed robust increases or decreases in fluorescence in response to thermal stimulation of the hindpaw (215 responsive / 2221 expressing neurons in *n* = 12 awake mice) (**Fig 1b,c, Supplementary Data Fig 1a-d**). We identified 3 major response profiles (**Fig 1d**): (i) neurons with an increase in fluorescence for cooling (‘Cool-ON’, 73%), (ii) neurons with an increase in fluorescence for warming (‘Warm-ON’, 7%), and (iii) bidirectionally modulated neurons with an increase in fluorescence for cooling and a reduction in fluorescence for warming (‘Cool-ON/Warm-OFF’, 18%). Occasionally, we observed rare response types (‘Other’ in **Fig 1**), including an increase in fluorescence for both warming and cooling (‘Warm-ON/Cool-ON’, 1%), as well as neurons with an increase in fluorescence for warming and a decrease for cooling (‘Warm-ON/Cool-OFF’, 1%) (**Supplementary Data Fig. 2**).

We observed that neurons with an enhanced or ‘ON’ response to cooling dominated the population representation (**Fig 1e, left**), closely reflecting the larger proportion of cells responsive to cooling than warming at downstream stages of the thermosensory pathway^14,15,35^. Cool-ON/Warm-OFF neurons made up a slightly smaller proportion of cooling response neurons (20%), but showed larger cool response amplitudes than Cool-ON neurons (**Fig 1f**) with similar latencies (**Fig 1g**). Notably, the most common response to warming was the reduced or ‘OFF’ response observed in the bidirectional Cool-ON/Warm-OFF population (**Fig 1e**, right, 65%), suggesting that this is central to the encoding of warm. In support, Cool-ON/Warm-OFF neurons had a shorter latency response to warm as compared to the Warm-ON neurons (**Fig 1g**), suggesting a higher sensitivity to warm.

### Anesthesia does not alter thermoreceptor encoding of temperature

The use of anesthesia during in vivo recordings offers a more stable recording condition, however it could alter neuronal responses to thermal stimuli^36,37^. To determine the impact of anesthesia on innocuous temperature encoding by thermoreceptors, we compared responses of the same cells during isoflurane anesthesia with their responses during wakefulness (**Fig 2a,b, Supplementary Data Fig 3a-c**). Overall, we observed highly similar responses to hindpaw cooling (22°C) or warming (42°C) in individual thermoreceptors under both conditions (**Fig 2a,b**). Across the population, we observed that all neurons with a thermal response under anesthesia maintained their characteristic functional tuning in awake mice (**Fig 2c**). However, some neurons (23%) with detectable responses in awake mice showed no significant response under anesthesia (**Fig 2d**). These neurons were characterized by a weak thermal response in awake mice rather than any specific response property (**Supplementary Data Fig 3d**). Overall, the proportion of the major functional response types (Cool-ON, Cool-ON/Warm-OFF, and Warm-ON) under anesthesia compared to awake mice was similar (**Fig 2e**). Moreover, neither functional response amplitudes nor response latencies were altered in thermoreceptors following anesthesia (**Fig 2f**). Given the similarity of responses in anesthetized and awake mice, we went on to perform imaging under anesthesia to allow a more in-depth investigation of thermoreceptor encoding properties.

**Figure 2.**
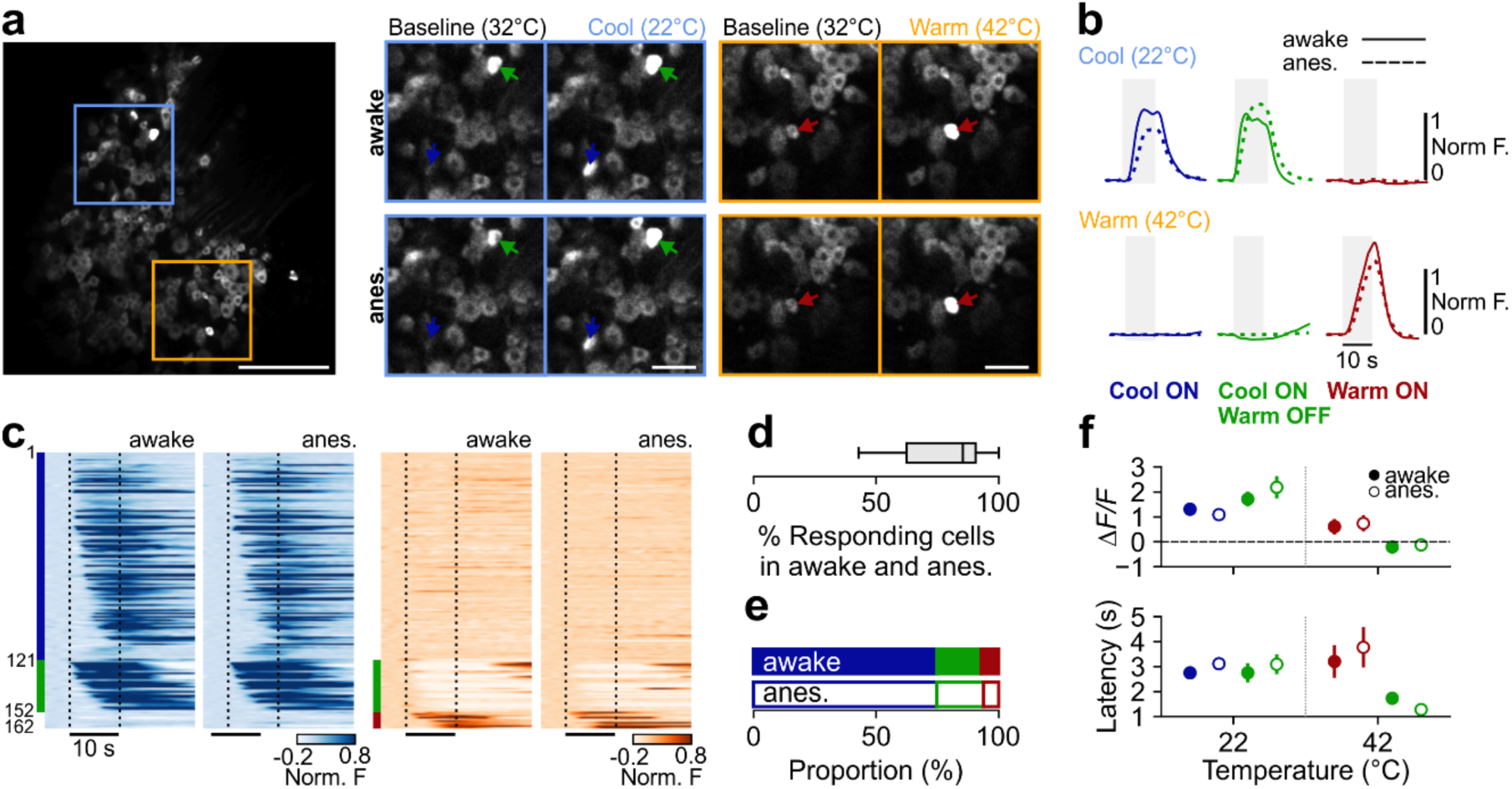
Thermoreceptors show the same thermal tuning in awake and anesthetized mice. **a.** Left, example *in vivo* two-photon image of TRPV1-GCaMP6s expressing neurons in L4 DRG. Right, the same example *in vivo* two-photon imaging plane in awake (top) and anesthetized (bottom) conditions at adapted temperature (AT) of 32°C, during a cooling stimulus (22°C), and during a warming stimulus (42°C). Left scale bar, 300 µm, right scale bar, 100 µm. Colored arrows show individual cells identified in both conditions, with color corresponding to response type as indicated in panel b. **b.** Example mean responses of single thermoreceptors in awake (solid line) or anesthetized (dashed line) conditions during a cooling (top) or warming (bottom) stimulus from an AT of 32°C (*n* = 10 trials each). Cell colors correspond to arrows depicted in a. Shaded gray box indicates length of stimulus (10 s). **c.** Single thermoreceptor responses to cooling (left, 22°C) and warming (right, 42°C) stimuli in awake and anesthetized mice. Each line represents a single neuron; sorted based on response latency and response type. Vertical dashed lines indicate the onset and end of the plateau phase of a thermal stimulus (*n =* 162 thermally responsive neurons, 12 mice). Index bar on the left indicates thermal responsivity for each neuron as indicated in panel b. **d.** Proportion of recorded thermoreceptors that are anatomically identified in both awake and anesthetized sessions and remain thermally responsive (mean 77%, from 12 mice). **e.** Proportion of thermally responsive sensory afferent neurons (*n =* 121 Cool-ON, blue; *n =* 31 Cool-ON/Warm-OFF, green; *n =* 10 Warm-ON, maroon, from 12 mice) in awake (closed circle) and anesthetized sessions (open circle). **f.** Top, Peak response amplitudes for cool (left) and warm (right) responses from each response classification as indicated in panel b for awake (closed circle) and anesthetized (open circle) sessions. Bottom, onset latencies for cool (left) and warm (right) responses from each response classification as indicated in panel b for awake (closed circle) and anesthetized (open circle) sessions. All data values show mean ± s.e.m..

### Thermoreceptors show graded encoding of cooling and warming amplitude

Two main encoding schemes have been proposed to describe how thermoreceptors might respond to different temperature amplitudes (**Fig 3a**): (i) as combinations of neurons that activate at specific temperatures ranges (specificity model), or (ii) as graded responses that monotonically change with increasing temperature (graded model). Warming has been described as following a graded encoding model, while the model for cooling remains unclear^17,29^. As with our data using Δ10°C stimuli in awake mice (**Fig 1**), only a small fraction of the GCaMP6 expressing population were responsive to any thermal stimuli (*n =* 499 out of 20,561 recorded neurons, *n =* 30 mice). Moreover, we did not observe any differences in the baseline fluorescence or cell size between responsive and unresponsive neurons (**Supplementary Fig 1f-i**). To determine the encoding strategy by thermoreceptors, we measured responses to different amplitudes of cooling or warming stimuli from an AT of 32°C (**Supplementary Fig 1e**) and went on to classify the thermal response profile of responsive neurons as consistent with either a specificity or a graded model (**Fig 3a-c**, see methods). More than 99% of the entire population of cool-responsive neurons (including Cool-ON, Cool-ON/Warm-OFF, and Warm-ON/Cool-ON) showed graded encoding to cooling (**Fig 3b-f**). Notably, we observed distinct tuning and recruitment profiles with respect to functional cell-type classification (**Fig 3d-f**). Cool-ON neurons showed a graded response profile that scaled linearly with cool amplitude (**Fig 3f**), with additional neurons gradually recruited as stimulus amplitude increased up to -Δ16°C (**Fig 3g**). Cool-ON/Warm-OFF neurons were more sensitive (**Fig 3f**), and were almost all recruited (show a significant change in fluorescence) at small amplitudes (-Δ2°C, **Fig 3g**).

**Figure 3.**
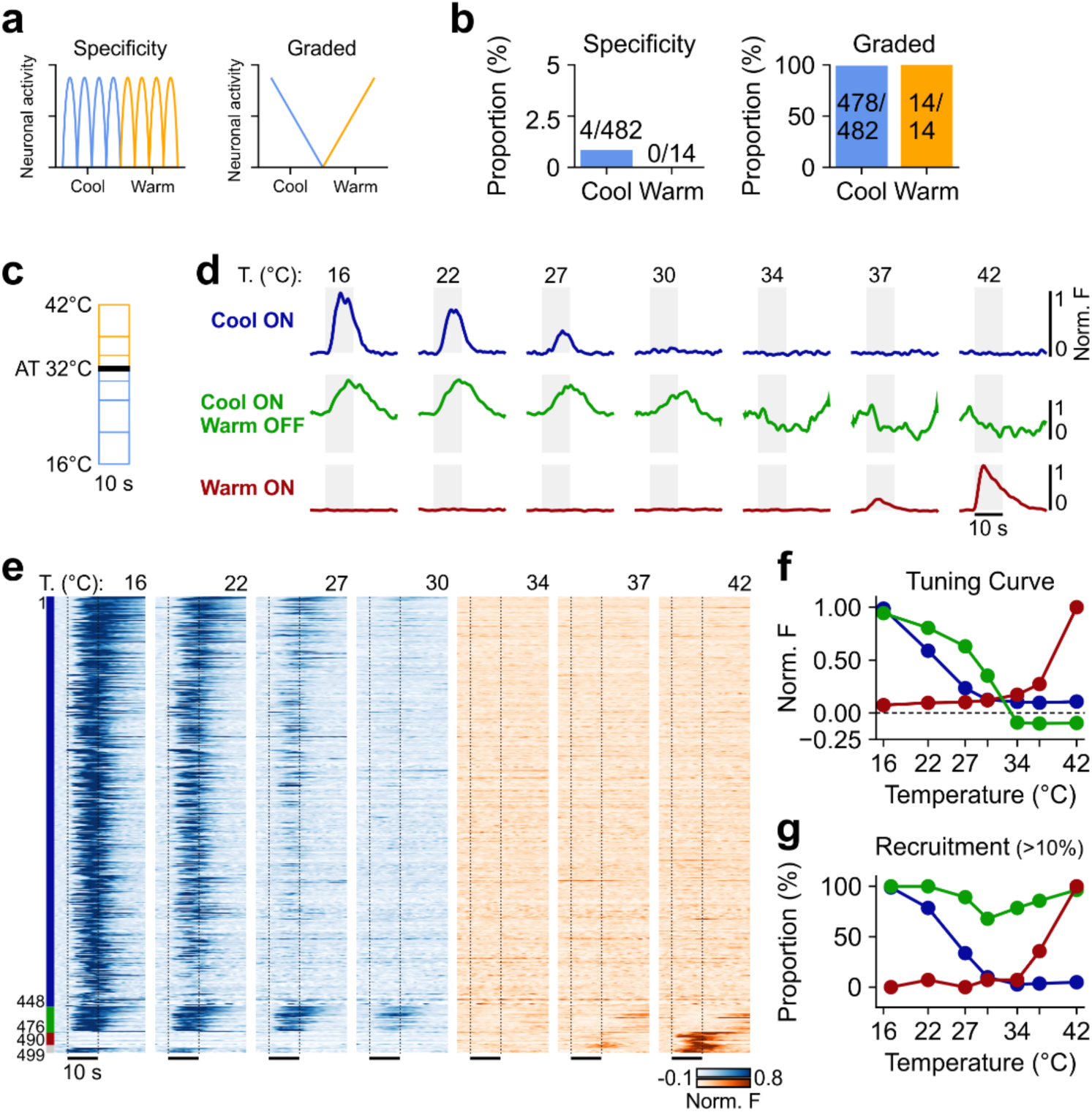
Thermoreceptor encoding of cool and warm amplitude. **a.** Schematic showing two possible encoding schemes for cool and warm. **b.** Proportion of neurons that follow a specificity encoding (left) or graded encoding (right) in response to cooling (blue, 16°C) or warming (orange, 42°C) **c.** Schematic of stimulus protocol used to test the sensitivity of thermal encoding. **d.** Trial average (*n =* 3 trials per temperature) traces of example thermoreceptors responding to cooling and warming from AT of 32°C. **e.** Single-cell thermoreceptor calcium responses to cooling (left – blue) and warming (right – orange) stimuli. Each line represents a single neuron; sorted based on both the response duration and the response type. Vertical dashed lines indicate the onset and end of the plateau phase of a thermal stimulus (*n =* 499 thermally responsive neurons, 30 mice). **f.** Normalized stimulus-evoked fluorescence across target temperatures for Cool-ON (blue; *n =* 448 neurons), Cool-ON/Warm-OFF (green; *n =* 28 neurons), and Warm-ON (maroon; *n =* 14 neurons). **g.** Proportion of neurons recruited (significant increase or decrease in fluorescence) to at least 10% of the maximum fluorescence response across target temperatures. *n* values as in f. Values in f and g show mean ± s.e.m..

All warm-responsive neurons showed graded response profiles to innocuous warming stimuli (**Fig 3b-f**), with more neurons recruited as warming stimuli increased in amplitude (**Fig 3g**). The Cool-ON/Warm-OFF population of neurons did not show an obvious graded reduction of activity to increased warming stimuli at an AT of 32°C (**Fig 3d-f**), with the majority of Cool-ON/Warm-OFF neurons recruited by small amplitude warming (+Δ2°C, **Fig 3g**), similar to their higher sensitivity for small amplitude cooling. Together these data highlight the bidirectional Cool-ON/Warm-OFF population of thermoreceptors as the most sensitive encoders of innocuous temperature changes, suggesting that they may constitute the main functional cell type for encoding both cool and warm.

### The adapted temperature reveals thermoreceptor functional subtypes

Two primary parameters define the thermal stimulus amplitude: the target temperature (TT), and the adapted temperature (AT). Thus far, we have considered the impact of varying the TT from a fixed AT of 32°C. However, while 32°C is considered a standard AT in humans and primates, neutral skin temperature varies significantly across body parts^38^ and mouse forepaw temperature has been estimated to be lower than 32°C (∼24-27°C, see ^7^). Further, a lower AT leads to greater acuity for warmth perception in mice^7^, suggesting that lowering the AT may result in recruitment of a larger and/or more sensitive population of Warm-ON thermoreceptors.

To address this hypothesis, we investigated the effect of AT on thermoreceptor encoding. Sensory evoked responses were measured to repeated cooling and warming TT from four ATs (22, 27, 32, 37°C, **Fig 4a**). We observed robust cellular responses to cooling and warming at all ATs and identified the different functional response groups we had observed from an AT of 32°C (**Fig 4b,c**). Moreover, applying the functional classification at each baseline, we saw that changing the AT significantly altered the relative proportion of functional subgroups (**Fig 4d**). At lower ATs, there was no increase in thermoreceptors excited by warm, rather, the proportion of Warm-ON and Warm-ON/Cool-OFF populations remained sparse and represented < 2% of the total thermosensory population (**Fig 4d**). Intriguingly, lower ATs resulted in a significant increase in the prevalence of Cool-ON/Warm-OFF thermoreceptors (**Fig 4d**). Together these data highlight that at lower ATs, the population response to warm is dominated by a reduction in the activity of Cool-ON/Warm-OFF thermoreceptors.

**Figure 4.**
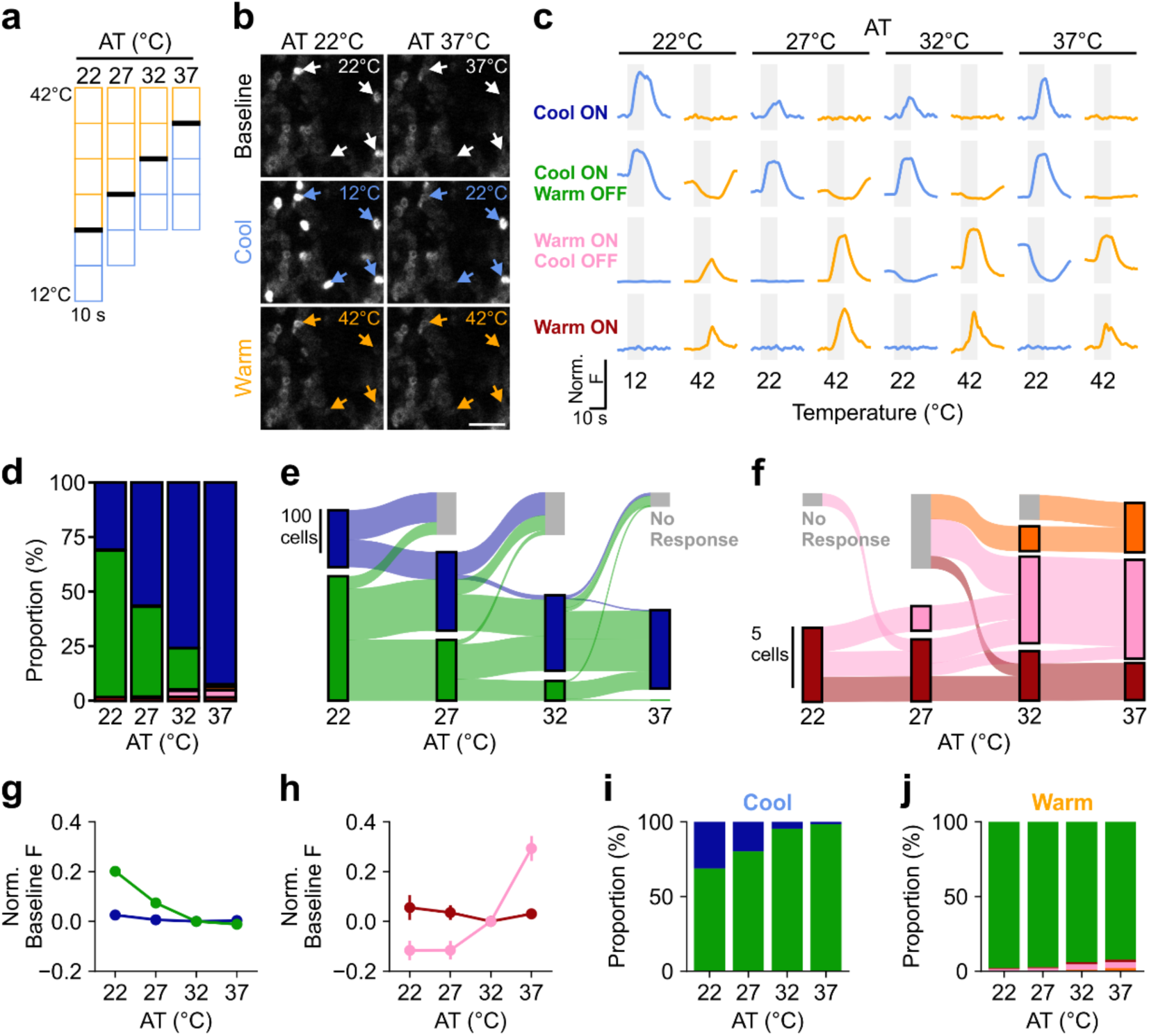
Changing the adapted temperature uncovers thermoreceptor functional response type. **a.** Schematic of stimulus protocols. **b.** Example of *in vivo* two-photon images at an AT of 22°C (left) and AT of 37°C (right) during baseline (top), a cooling (middle 12°C or 22°C), and a warming stimulus (bottom 42°C). Scale bar, 100 µm. **c.** Trial average (*n =* 3 trials per temperature) traces of example thermoreceptor response to cooling and warming from AT of 22, 27, 32, and 37°C. **d.** Proportion of thermally responsive afferent neurons at different ATs (AT 22°C, *n =* 416 neurons; AT 27°C, *n =* 353 neurons; AT 32°C, *n =* 262 neurons; AT 37°C, *n =* 250 neurons, 16 mice). **e.** Sankey plot depicting transitions of thermoreceptors between functionally defined Cool-ON thermosensory response profiles over all AT. Responsive cells are classified according to thermal tuning preference, with neurons classified as none if no temperature evoked calcium transients were observed. Bar color indicates classification at each AT while ribbon color indicates classification at the lowest AT (22°C). **f.** Same as e except for functionally defined Warm-ON thermosensory response profiles with ribbon color indicating classification at the highest AT (37°C). **g.** Normalized baseline fluorescence values across AT for response classifications with an ON response to cooling (*n =* 124 Cool-ON, blue; *n =* 280 Cool-ON/Warm-OFF, green). **h.** Same as g except for neurons with an ON response to warming (*n =* 5 Warm-ON, maroon; *n =* 8 Warm-ON/Cool-OFF, pink). **i.** Proportion of cool responsive neurons across AT with neuron populations classified at the lowest AT (22°C). **j.** Same as i except for warm responsive neurons classified at highest AT (37°C). Cell type colors follow the same scheme as written in c, data points show mean ± s.e.m..

Next, we sought to examine how functional identity changes across ATs for individual thermoreceptors. We found that OFF responses were most prominent at lower ATs for Cool-ON/Warm-OFF thermoreceptors, and at higher ATs for Warm-ON/Cool-OFF thermoreceptors (**Fig 4d**). We therefore asked whether thermoreceptors classified as ‘ON only’ at one AT could have an OFF response unmasked by changing AT. To visualize and compare functional response classification across AT for individual thermoreceptors, we constructed Sankey plots that tracked the functional response type defined at each baseline whilst maintaining the response type classification at the lowest (**Fig 4e**) or highest (**Fig 4f**) AT (bar color shows cell response type redefined at each AT as in **Fig 4d**, while ribbon color shows response type defined at an AT of 22°C or 37°C for **Fig 4e** or **4f** respectively). We found that almost all thermoreceptors with an ON response to cooling at 37°C AT (**Fig 4e**, blue bar at AT 37°C) had an OFF response progressively unmasked with decreasing AT (**Fig 4e**, green ribbon extending from AT 37°C). A similar change in functional response classification with AT was observed in the sparse population of neurons with ON responses to warming (**Fig 4f**).

Therefore, across neurons with an excitatory response to thermal stimulation, changes in the AT did not alter the excitatory selectivity. Instead, changes in the AT only change the presence of an OFF response. Because we recorded from each neuron in every stimulus condition, we were able to redefine the functional response classification system across AT to assign a single functional classification for each cell. If the cell had the capacity to display an OFF response at any AT, it was defined as an ON/OFF classification. If no OFF response was identified at any AT, it was defined as an ON classification. Using this re-classification, we first investigated how an OFF response can be unmasked by AT. Across the population of recorded neurons, we observed that Cool-ON/Warm-OFF thermoreceptors showed higher baseline fluorescence prior to stimulus onset at the lowest AT (22°C) (**Fig 4g**, example in **Fig 4b, top**). This was not observed in Cool-ON thermoreceptors, whose baseline fluorescence remained unchanged regardless of AT (**Fig 4g**). Similarly, Warm-ON/Cool-OFF thermoreceptors had higher baseline fluorescence at the highest AT (37°C, **Fig 4h**), whilst the baseline fluorescence of Warm-ON thermoreceptors remained unchanged (**Fig 4h**). This suggests that the OFF response is enabled by the presence of ongoing baseline activity at different ATs. Further, these data reveal that the Cool-ON/Warm-OFF thermoreceptors, re-defined across baselines, are by far the largest group of thermoreceptors that encode both innocuous cooling (**Fig 4i**) and warming (**Fig 4j**) and therefore are the critical neuronal population for innocuous thermosensation.

### Thermoreceptors encode absolute temperature changes

As with all sensory systems, and observed in **Fig 4**, responses to thermal stimuli vary with respect to the background sensory input. In the thermal system, it has been proposed that thermoreceptors show warming responses that reflect the absolute temperatures value of the stimulus (an absolute encoding scheme), while cooling responses reflect the difference in amplitude between the stimulus and adapted temperature value (a relative encoding scheme)^39^. However, to date there has never been a systematic investigation of absolute and relative encoding by the same thermoreceptors to innocuous thermal stimuli.

If a neuron follows a relative encoding scheme, there should be no change in response amplitude across different ATs when the difference between the AT and stimulus amplitude is fixed (**Fig 5a**). We first tested thermoreceptor response amplitude to changes in temperature (ΔT) of the same amplitude from different ATs. It is evident from the responses of different functional cell types to an example fixed ΔT for cooling (**Fig 5b** (-10°C), **Supplementary Data Fig 4a** (-5°C), blue) or warming (**Fig 5b** (+5°C), **Supplementary Data Fig 4a** (+10°C), orange) that the response strength varied across ATs, indicating that thermoreceptors do not follow a relative encoding scheme for cool or warm. A difference in response amplitudes for a fixed ΔT can also be seen in the subpopulation mean responses at different ATs (**Supplementary Data Fig 4c**). If thermoreceptor activity instead reflects the absolute temperature values of a stimulus, there should be no change in the response amplitude when presenting the same absolute target temperature from different ATs (**Fig 5c**, TT). Visualizing the thermoreceptor responses to an example fixed TT (22°C, **Fig 5d**, blue; or 42°C, **Fig 5d**, orange) from different ATs, and plotting the mean subpopulation response amplitudes against TT (**Supplementary Data Fig 4b**), showed similar response amplitudes to different TT across ATs, consistent with an absolute encoding scheme.

**Figure 5.**
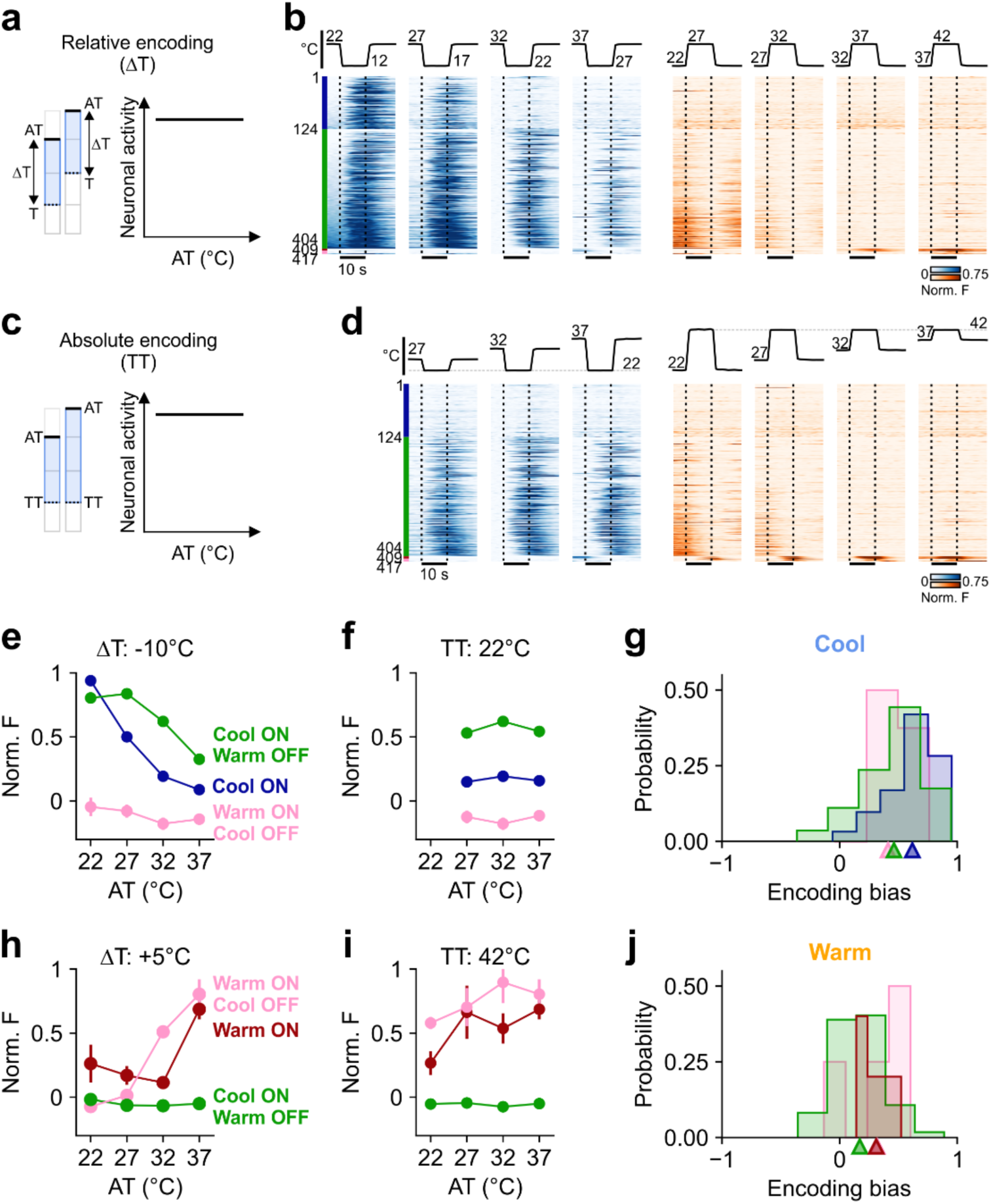
Thermoreceptors encode absolute temperature changes. **a.** Schematic of relative encoding scheme whereby neural activity is dependent on the change in stimulus temperature (ΔT) relative to the adapted temperature (AT). **b.** Single thermoreceptor responses to cooling (ΔT = -10°C, left – blue) and warming (ΔT = +5°C, right – orange) stimuli from different AT. Each line represents a single neuron; sorted based on both the baseline fluorescence in the lowest AT (22°C), and the response type. Vertical dashed lines indicate the onset and end of the plateau phase of a thermal stimulus (*n =* 417 thermally responsive neurons, 16 mice). **c.** Schematic of the absolute encoding scheme whereby neural activity is dependent on the target stimulus temperature (TT) independent of the adapted temperature (AT). **d.** Same as b, but for a fixed cooling (TT = 22°C) and warming (TT = 42°C) target temperature. **e.** Normalized peak fluorescence response values across AT in response to a ΔT of -10°C for response classifications with an ON (*n =* 124 Cool-ON, blue; *n =* 280 Cool-ON/Warm-OFF, green) or OFF response to cooling (*n =* 8 Warm-ON/Cool-OFF, pink). **f.** Same as e., but for TT = 22°C. **g.** Encoding bias, defined as the similarity of an individual unit tuning to absolute (+1) or relative (-1) encoding in response to cooling. Colors indicate functional response classification. **h.** Same as e except for ΔT = +5°C for response classifications with an ON (*n =* 5 Warm-ON, maroon; *n =* 8 Warm-ON/Cool-OFF, pink) or OFF response to warming (*n =* 280 Cool-ON/Warm-OFF, green). **i.** Same as i except for TT = 42°C. **j.** Encoding bias for warming responses. Colors indicate functional response classification as depicted in e and h, and all data values depict mean ± s.e.m.

To further analyze the cooling data, we plotted the population mean responses of each functional subgroup to a fixed ΔT or a fixed TT cooling stimulus against the 4 ATs (**Fig 5e,f**). As observed in the individual thermoreceptor responses, and inconsistent with a relative encoding scheme for cooling, the mean response amplitude to a fixed ΔT decreases with increasing AT (**Fig 5e, Supplementary Data Fig 5a**). If instead we consider a fixed cooling TT, the average response amplitude is approximately equivalent across different ATs (**Fig 5f, Supplementary Data Fig 5b**), consistent with an absolute encoding scheme. Across all ATs, we noted that Cool-ON/Warm-OFF thermoreceptors show a higher sensitivity to cooling than Cool-ON thermoreceptors (**Fig 5e,f, Supplementary Data Fig 5a,b**). Grouping Cool-ON/Warm-OFF neurons by the first AT at which an OFF response was observed (**Supplementary Data Fig 6a**) also demonstrates absolute encoding across sensitivity levels (**Supplementary Data Fig 6b,c**).

To quantify the degree of relative or absolute encoding for individual thermoreceptor to cooling, we computed an encoding bias parameter (see Methods) across all stimulus conditions with more than one stimulus per AT (**Fig 4a**). This parameter is -1 if the neuron is a perfect relative encoder (identical response amplitude for every fixed ΔT across all AT) or is +1 if the neuron is a perfect absolute encoder (identical response amplitude for every fixed TT across all AT). The distribution of cooling responsive neurons was strongly biased towards absolute encoding (**Fig 5g, Supplementary Data Fig 6d**).

Similar analysis of thermoreceptor responses to warming, also showed that the TT was a more reliable predictor of response amplitude compared to the change in temperature (ΔT) when plotted against the AT, consistent with an absolute encoding of warming (**Fig 5h,i,j**; **Supplementary Data Fig 5c,d**). Moreover, OFF responses for warming and cooling had very little dynamic range but similar analysis support absolute encoding (**Supplementary Data Fig 5e,f**). Together, our data reveal that all functional subtypes of thermoreceptors represent absolute changes in temperature both for cool and warm.

### Thermoreceptor encoding of temperature is dependent on TRPM8

TRPM8 is a temperature sensitive ion channel expressed in thermoreceptors that is selective for cool. However, TRPM8^-/-^ mice have a deficit in the perception of both cool and warm^7,22^, suggesting this channel might be involved in Cool-ON/Warm-OFF responses. Given the prevalence and sensitivity of Cool-ON/Warm-OFF responses, we asked whether TRPM8 could be required for both the enhanced response to cooling as well as the reduced response to warming.

We first assessed the contribution of TRPM8 to cold-responding thermoreceptor activity (Cool-ON and Cool-ON/Warm-OFF) using *in vivo* pharmacology to block the channel’s activity. The TRPM8 selective antagonist PBMC (Phenylethyl-(2-aminoethyl) [4-(benzyloxy)-3-methoxybenzyl] carbamate^40^) was injected into the plantar surface of the hindpaw. Thermal responses were measured immediately prior to injection and 15 min afterwards (**Fig 6a**). Application of PBMC caused a complete loss of cool-evoked responses in both Cool-ON and Cool-ON/Warm-OFF neurons across ATs (**Fig 6c**). Inhibition of TRPM8 also resulted in a marked decrease of baseline fluorescence in Cool-ON/Warm-OFF neurons at an AT of 22°C (**Fig 6b**). The loss of ongoing neural activity, as indicated by baseline fluorescence, also eliminated the warm suppressed responses in Cool-ON/Warm-OFF neurons following PBMC administration, consistent with a lack of Cool-ON/Warm-OFF cells in TRPM8^-/-^ mice ^7^.

**Figure 6.**
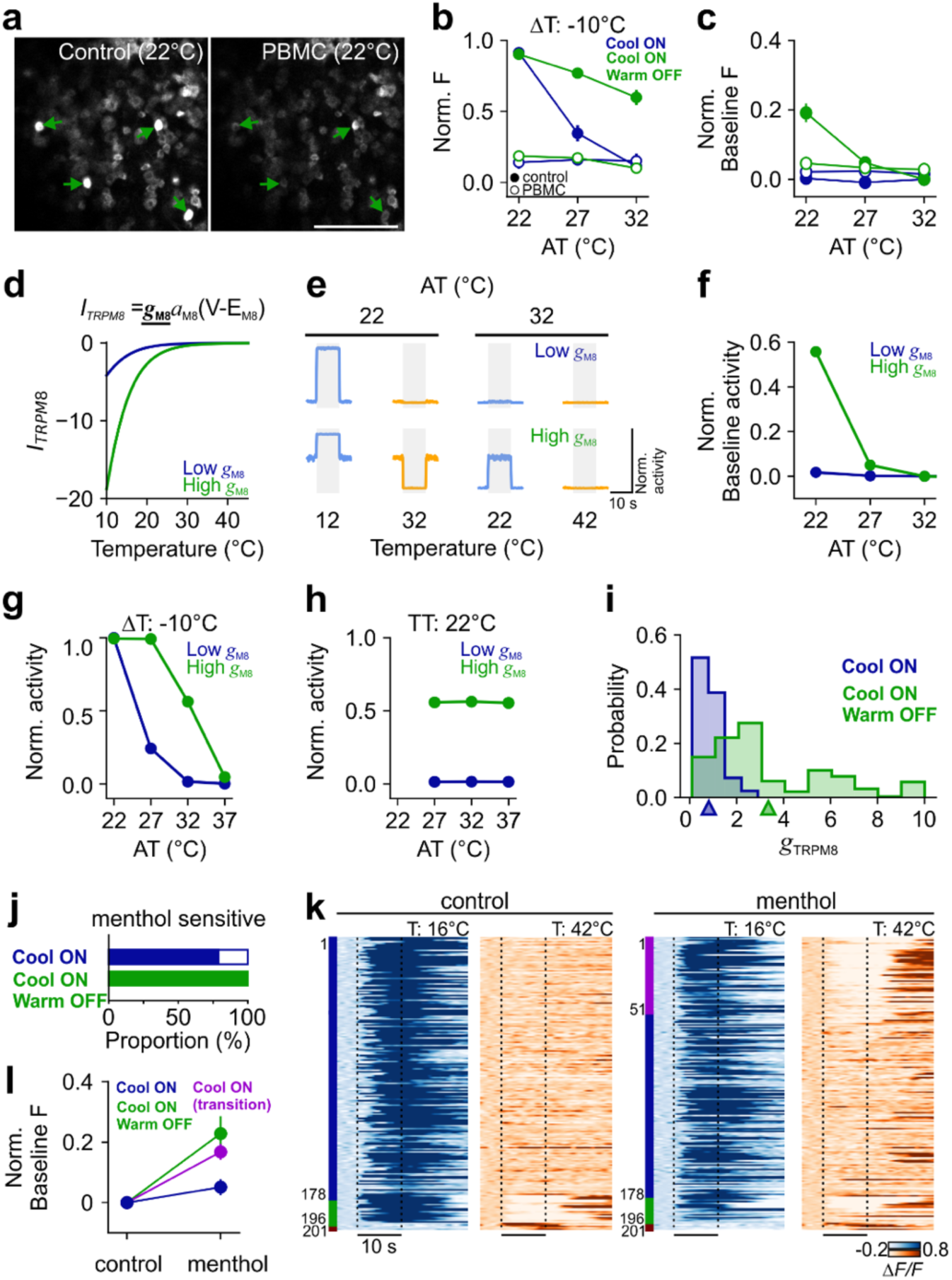
Cool-ON and Warm-OFF responses are mediated by TRPM8. **a.** Example *in vivo* two-photon image of thermoreceptors at an AT of 22°C before (left) and after (right) application of TRPM8 antagonist, PBMC. Responsive neuron ROIs are denoted by arrows colored according to functional classification (Cool-ON/Warm-OFF, green), Scale bar, 300 µm. **b.** Normalized baseline fluorescence values across AT for response classifications with an ON response to cooling before (solid marker) and after (open marker) application of PBMC (*n =* 40 Cool-ON, blue; *n =* 41 Cool-ON/Warm-OFF, green). **c.** Normalized peak stimulus-evoked fluorescence response values across AT in response to a ΔT of -10°C for the cells presented in b. **d.** Top, equation for a cool-responsive current (I_TRPM8_). Bottom, modelled TRPM8 current flow for a low (g_M8_=0.5 mS/cm^2^, blue) and a high (g_M8_=2.25 mS/cm^2^, green) conductance value at a fixed membrane voltage (V_m_ = -30mV). **e.** Simulated model response to cooling (blue, -10°C) or warming (orange, +10°C) stimuli across AT (22°C and 32°C) for low (top) and high (bottom) conductance values. **f.** Simulated normalized baseline firing rate values across AT for low (blue) and high (green) conductance values. **g.** Model simulation normalized firing rate response values across AT in response to a ΔT of -10°C as depicted in the stimulus trace schematics in Figure 4c for low (blue) and high (green) conductance values. **h.** Same as g except for TT = 22°C. **i.** Model-fit TRPM8 conductance values to each experimental neuron with a Cool-ON response (*n =* 184 Cool-ON, blue; *n =* 280 Cool-ON/Warm-OFF, green). **j.** Proportion of neurons that display a response to the application of TRPM8 agonist, menthol, to the skin by response classification (blue: Cool-ON, *n =* 178 neurons; green: Cool-ON/ Warm-OFF, *n =* 18 neurons). **k.** Single thermoreceptor calcium responses to cooling (AT: 32°C, TT: 16°C, blue) and warming (AT: 32°C, TT: 42°C, orange) stimuli before (left) and after (right) application of menthol. Each line represents a single neuron; sorted based on both the response width, and the response type. Vertical dashed lines indicate the onset and end of the plateau phase of a thermal stimulus (*n =* 201 thermally responsive neurons, 7 mice). **l.** Baseline fluorescence before and after application of menthol for cells classified as Cool-ON/Warm-OFF in control conditions (green, *n =* 18 neurons) and for cells classified as Cool-ON cells in control conditions that transition to Cool-ON/Warm-OFF with menthol application (magenta, *n =* 51 neurons) or that do not transition with menthol application (blue, *n =* 127 neurons). All data values depict

Given the absence of background activity at low ATs during pharmacological blockade of TRPM8, we hypothesized that higher levels of TRPM8 conductance could modulate baseline activity at different ATs and, following channel inactivation to warm, suppress their activity, while also signaling cool by an increase in activity. To address this hypothesis, we generated a leaky integrate-and-fire neuron model with a cool-responsive TRPM8 current^41,42^. The modelled TRPM8 current retained the known channel characteristics^43^ but allowed the maximum conductance (gM8) to vary within physiological limits. Low conductance model neurons (0.5 mS/cm^2^) have lower TRPM8 current contributions than high conductance (2.25 mS/cm^2^) model neurons (**Fig 6d**). The low gM8 model neuron displayed the response profile seen experimentally in Cool-ON neurons at different ATs (see **Fig 4c**), where thermoreceptors respond robustly to large cooling stimuli, weakly to low cooling stimuli, and have no response to warming (**Fig 6e**). The high gM8 model neuron mimicked responses seen experimentally in Cool-ON/Warm-OFF thermoreceptors (see **Fig 4c**), where cooling-evoked activity was more sensitive and a robust reduction in activity was seen to warming due to the increased baseline activity (**Fig 6e,f**). Further, the model closely aligns with the experimental data (**Fig 5e,f**) such that the model neuron followed an absolute encoding scheme to cooling stimuli (**Fig 6g,h**) for both the low and high gM8 conditions. Therefore, varying only the TRPM8 conductance enables the generation of the two distinct cool response profiles seen experimentally.

As such, the model predicts that the primary difference between Cool-ON and Cool-ON/Warm-OFF thermoreceptors is their level of TRPM8 conductance. Biologically this could be equivalent to differing expression levels of TRPM8. Model neuronal responses were generated for gM8 values ranging from 0.1 to 10 mS/cm^2^ and compared to experimentally recorded neuronal responses to estimate a best fit gM8 value for each experimental neuron. TRPM8 conductance values estimated for experimentally recorded neurons were higher in Cool-ON/Warm-OFF than Cool-ON thermoreceptors (**Fig 6i**), consistent with the model hypothesis that TRPM8 conductance defines the differentiation of these two functional groups. Estimated TRPM8 conductance values were also highest for higher sensitivity Cool-ON/Warm-OFF thermoreceptors (**Supplementary Data Fig 6e**), further implicating TRPM8 conductance as a defining feature of the thermal response profile.

If TRPM8 conductance is the fundamental difference between Cool-ON and Cool-ON/Warm-OFF thermoreceptors, we would predict that artificially increasing the TRPM8 conductance could transform Cool-ON thermoreceptor response into Cool-ON/Warm-OFF responses. We tested this experimentally by increasing TRPM8 conductance with application of the TRPM8 agonist menthol^18^ to the hindpaw. Almost 75% of Cool-ON, and 100% of Cool-ON/Warm-OFF neurons were sensitive to menthol and, as predicted by the model, a proportion of menthol-sensitive Cool-ON neurons (*n* = 51 out of 178 neurons) showed an unmasking of warm evoked suppression following application of menthol (**Fig 6j, k**, magenta). The unmasked OFF response to warm was enabled by an increase in baseline fluorescence that was shifted to a level comparable to the Cool-ON/Warm-OFF population (**Fig 6l**, green). The thermoreceptors that did not transition from Cool-ON to Cool-ON/Warm-OFF with the application of menthol did not show a substantial increase in baseline fluorescence (**Fig 6l**, blue). These data indicate that simply increasing the level of TRPM8 conductance can result in Cool-ON thermoreceptors converting to Cool-ON/Warm-OFF thermoreceptors. This supports our findings using lower ATs (**Fig 3**) and suggests that possibly all Cool-ON neurons have the capacity to show a Warm-OFF responses. We conclude that TRPM8 is required for Cool-ON and Warm-OFF, accounting for almost all our innocuous thermal responses, and therefore is the key thermally sensitive channel for innocuous thermosensation.

## Discussion

To address how thermoreceptors encode innocuous cool and warm, we developed a chronic *in vivo* DRG calcium imaging preparation for awake and anesthetized mice. By presenting precise thermal stimuli encompassing a broad stimulus space, we examined competing thermal encoding models and went on to test the role of TRPM8 in thermal responses using computational modeling and *in vivo* pharmacology. Here we show that the majority of thermoreceptors report cool and warm bidirectionally with an increase or decrease in activity. Intriguingly, these bidirectional responses can both be attributed to changes in TRPM8 conductance.

Innocuous cool stimuli evoked robust activity in two major populations of thermoreceptors, warm-insensitive cool-only thermoreceptors (Cool-ON) and cool/warm sensitive thermoreceptors (Cool-ON/Warm-OFF). Cool-ON thermoreceptors were most strongly activated by large amplitude cooling stimuli and only showed recruitment of responsive cells as amplitudes approached noxious cooling. In contrast, Cool-ON/Warm-OFF thermoreceptors were more sensitive and more rapidly recruited at lower amplitude temperature values, indicating a role in innocuous thermal perception. Functional subpopulations of cool responsive thermoreceptors have also been determined by their cool threshold, with low and high cool thresholds broadly reported in previous work^17,28,32,44–46^. One possibility is that the expression level of TRPM8 dictates this difference in threshold^17,32,44^. In support, our pharmacological manipulations and computational modelling revealed how differences in TRPM8 conductance level can explain thermoreceptor sensitivity to cool (Figure 6). Differences in TRPM8 conductance are also able to define whether a cool responsive cell is Cool-ON or Cool-ON/Warm-OFF, and reveals that the majority of thermoreceptors that encode cooling are capable of a Warm-OFF response.

In agreement with prior work, we observed very few thermoreceptors with excitatory responses to warm^17,29,47,48^. Instead, we observed that innocuous warm is largely represented by a reduction of activity in Cool-ON/Warm-OFF thermoreceptors. The large numbers of Cool-ON/Warm-OFF cells and the high sensitivity of their response to warm, suggests that warm suppression is a key aspect of the encoding of low amplitude innocuous warm temperatures. In support, lowering the AT, which increases the proportion of Cool-ON/Warm-OFF thermoreceptors (Figure 4), has been shown to increase the acuity for warm perception in mice^7^. Moreover, elimination of TRPM8 leads to a pronounced deficit in the perception of innocuous warm and a notable reduction in the numbers of Cool-ON/Warm-OFF thermoreceptors^7,22,46^. Excitatory thermoreceptor responses to warm could be supported by other TRP channels, like TRPV1 or TRPM2^17,48,49^. These neurons could still contribute to the perception of warm, indeed there are deficits in warm perception in TRPV1^-/-^ and TRPM2^-/-^ mice^7,17^, but their major role might be in the perception of larger amplitude warm/hot stimuli or in the discrimination of different levels of warmth.

Behavioral studies have shown that cool and warm sensations show significant differences despite being on the same sensory axis. Warm has a higher perceptual threshold and is less accurately detected than cool by humans and mice^5–7^. Moreover, in the thalamus and cortex, warm neuronal responses are characterized by slower kinetics and higher activation thresholds compared to cool^14,15^. Alongside a smaller number of warm excitatory responses in thermoreceptors, another possible explanation for these differences is that central warm encoding relies on additional processing steps compared to cold. We show above that many thermoreceptors report warming with a reduction in activity, but prior work has shown that downstream neurons encode warm primarily with increases in activity^14,15,35,50^. Because excitatory responses to warming are already seen in spinal cord neurons^35,50^, one hypothesis is that inversion of warm signaling from suppression to excitation occurs in the spinal cord via a disinhibitory mechanism, as reported in first order thermal projection neurons in Drosophila^51^ and could provide a mechanistic understanding for the distinct encoding profiles of cool and warm seen in thermoreceptors. Future work to address this hypothesis will require recordings and manipulations of identified spinal cord neurons.

Temperature, like other sensory modalities, is encoded on top of a baseline of constant sensory input. In the skin, surface temperature is defined by a combination of external temperature and the animal’s own body temperature. Here we experimentally controlled the skin temperature by holding the Peltier element at specific adapted temperatures (ATs) before delivering brief thermal stimuli to test the contribution of relative and absolute encoding within the same thermoreceptors. Prior work has looked at the impact of AT on neuronal encoding of thermal stimuli at different stages of the thermal system, and suggested that warm stimuli are represented as their absolute stimulus level whereas cool by the change relative to the AT^8,14,29,35^. In agreement with prior work, our data show that warm is represented as an absolute encoding scheme. In contrast to prior work, responses to our broad stimulus set indicates an absolute encoding scheme for cool thermoreceptor responses, as recently observed in Drosophila thermoreceptors^52^. Moreover, incorporating a TRPM8 channel with fixed thermal sensitivity in our biophysical model of thermoreceptors recapitulates our experimental findings. Together we show that the afferent drive to the thermal system for innocuous temperature is driven by a representation of the absolute temperature for both cool and warm.

Our study represents the most comprehensive examination of innocuous thermal encoding by thermoreceptors *in vivo*. We show that the majority of thermoreceptors are bidirectional, encoding temperature with an increase in activity to cool and a decrease to warm. Notably, this bidirectionality is determined by the same thermal channel, TRPM8, and therefore provides support for the role of TRPM8 in warm perception. Our findings will be required for an understanding of how temperature is encoded at higher levels of the thermal system, and suggests that the same population of afferent neurons can be involved in generating highly distinct sensations.

## Funding sources

European Research Council ERC-2015-CoG-682422 (JFAP), Deutsche Forschungsgemeinschaft FOR 2143 (JFAP), Deutsche Forschungsgemeinschaft SFB 1315 (JFAP), National Institutes of Health NIH R01 NS123711 (JFAP), Helmholtz Association (JFAP), Human Frontiers Science Program LT000359/2018 (CJW).

## Competing interests

Authors declare that they have no competing interests.

## Data and materials availability

All data and code are available upon request.

We thank Charlene Wenzel, Nadine Gross, Svenja Steinfelder, Florian Rau for technical and administrative assistance, and all colleagues that read the manuscript.

## Contributions

PB, CJW, and JFAP designed the study. PB, CJW performed experiments and analyzed the data. PB, CJW, and JFAP wrote the manuscript.

## Methods

### Mice

All experiments were approved by the Berlin Landesamt für Gesundheit und Soziales (LAGeSo) and carried out in accordance with European animal welfare law. All mice were maintained on a 12:12 h light-dark cycle, with food and water provided *ad libitum* unless stated otherwise. Ai162D (B6.Cg-Igs7tm162.1(tetO-GCaMP6s,CAG-tTA2)Hze/J) and Ai148D (B6.Cg-Igs7tm148.1(tetO-GCaMP6f,CAG-tTA2)Hze/J) mice were crossed with TRPV1-cre^33^ mice to enable specific expression of GCaMP6s and GCaMP6f respectively in all thermosensory lineage neurons^33^ of the dorsal root ganglia (The Jackson Laboratory, stock no. #031562, #030328^53^). *TRPV1::Ai148D* (*n =* 31 mice) and *TRPV1::Ai162D* (*n =* 15 mice) adult female and male mice were used for calcium imaging experiments. All mice were housed and bred at the Max Delbrück Center for Molecular Medicine.

### Calcium imaging in dorsal root ganglia

#### Implantation of spinal clamp and chronic window for functional imaging

Mice were deeply anesthetized with ketamine (120 mgkg^-1^, i.p. beta-pharm) and xylazine (10 mgkg^-1^, i.p. Bayer), were administered with metamizol (200 mgkg^-1^, s.c. Zentiva) for peri- and postoperative pain management, and were secured in a stereotaxic frame (Narishige). Eye gel (Visc-Opthal, Bausc + Lomb) was applied to both eyes and core temperature was maintained at 37°C by a homeothermic heating blanket and rectal probe (FHC). The spinal clamp and DRG window were implanted as previously described in ^34^. Briefly, the L3-L5 vertebrae were exposed and cyanoacrylate glue and dental cement (Sun Medical C&B Super-Bond) were used to attach and fix the spinal clamp to the L3 and L5 vertebrae. Connective tissue overlying the neural foramen were dissected and removed to expose the L4 DRG. The DRG was covered with a thin layer of Kwik-Sil^®^ silicone elastomer (World Precision Instruments) and sealed using a 2 mm diameter glass window (Workshop of Photonics). Cyanoacrylate glue and dental cement were used to seal and fix the glass window to the vertebrae and spinal clamp. Mice that underwent acute experiments (*n =* 28 mice) were then transferred to the microscope for functional anesthetized imaging, as described below. Following an acute recording, mice were euthanized. Mice that underwent chronic experiments were transferred to a heated recovery chamber (38°C, Labotect) until ambulatory. Postoperative pain was managed by supplementing drinking water with metamizole (200 mgkg^-1^, Ratiopharm) for 3 days post-surgery.

#### Habituation to spinal clamp fixation and hindpaw restraint for awake functional imaging

For awake functional imaging, spinal clamp implanted mice were habituated to spinal clamp restraint in the imaging setup over a period of two weeks with increasing restraint times. During subsequent spinal habituation sessions, mice were also habituated to hindpaw restraint with increasing restraint times using medical tape (Leukosilk). Mice were regularly rewarded with sweetened condensed milk during the habituation process.

#### Imaging configuration for calcium imaging in dorsal root ganglia

Two-photon imaging was either performed acutely on the day of surgery (anesthetized), or chronically (anesthetized or awake) following the spinal clamp and DRG window implant. Fluorescence signals in anaesthetised (1.5% isoflurane in oxygen) and awake mice were recorded using a custom-built resonant-galvo two-photon microscope with a Nikon 16x water-immersion objective (0.8 NA, 3 mm WD, 1X FOV of 975 µm x 975 µm), equipped with a Chameleon Ultra II (Coherent) laser tuned to 920 nm (up to 80 mW power at the sample). The microscope was operated using ScanImage^®^ software (MBF Bioscience). Single plane acquisition was performed at an acquisition rate of 30 Hz and varying magnifications determined by the size of the DRG. Multiplane acquisition (20 µm spacing) was controlled by a fast piezo objective scanner (ThorLabs), with up to seven planes acquired sequentially (3.75Hz volume rate) dependent on the signal quality. Acquisition trials lasted 25 - 30 s and had an inter-trial interval of 80 s which was not recorded to minimize photobleaching. Stimulus generation, hardware synchronization, and data acquisition were performed using a National Instruments Data Acquisition card running custom-written Matlab code.

#### Sensory stimulation paradigms

Multiple thermal stimulation paradigms were administered to the hindpaw of the mice. Each stimulus set was presented once during a single recording session. A single mouse could undergo multiple stimulus conditions across multiple days. The entire surface of the glabrous skin of the right hindpaw was tethered onto the center of a Peltier-based thermal stimulator (QST-lab). Thermal stimulus sets were designed to investigate multiple aspects of thermal encoding with varying target temperatures (12°C to 42°C) and four adaptation temperature values (22, 27, 32, 37°C). In all conditions, thermal stimuli of 10 s duration were interleaved randomly every 100 s. The hindpaw was adapted to different adaptation temperatures (AT) over 80 s. The Peltier stimulator is held at this temperature until a transient stimulus is delivered (target temperature). Stimulus sets presented in awake mice were shortened to minimize the duration of restraint.

#### Pharmacological manipulations

A reduced thermal stimulus set was first recorded without pharmacological manipulation. Following administration of the substance, the imaging session was resumed following a 15 minute break using the identical stimulus set. Pharmacological manipulations were performed on separate days, separated by at least one week. TRPM8 agonist, menthol (640 mM), was applied topically to the plantar surface of the hindpaw. TRPM8 antagonist, PBMC (1-phenylethyl-4-(benzyloxy)-3-methoxybenzyl(2-aminoethyl)carbamate, 10 mg/ml diluted in 40% DMSO, Focus Biomolecules) was injected (∼10 µL, 0.3 ml insulin needle) transdermally into the glabrous skin of the right hindpaw in the anesthetized mouse.

#### Calcium imaging data analysis

Two photon imaging data were segmented anatomically into regions of interest (ROI) using a custom image segmentation model (CellPose,^54^) and functional responses were extracted using Suite2p^55^. ROIs that do not correspond to a cell body were manually excluded from further analysis. Movement in the image was assessed by the amount of required motion correction on each frame. Some trials were excluded if movement artefacts were too great and functional responses were not reliably extracted. ROIs were parameterized by size (cross sectional area), stability (fluorescence over the recording), and brightness (baseline fluorescence). Responsive ROIs were defined as stimulus-evoked fluorescence responses that were statistically different from the baseline fluorescence in the five seconds prior to stimulus onset. Responsive ROIs in multiplane recordings were spatially correlated across planes to control for optical contamination due to narrow z-plane spacing. ROIs assessed across multiple days or conditions were anatomically correlated across stimulus sets to control for variability in the recording plane. To do so, the average image for each condition was anatomically segmented into ROIs using the custom model (CellPose^54^). ROIs were only included in the longitudinal or multi-condition comparisons if the ROI was anatomically detected in both sessions and the overlap with the Suite2p identified ROI was greater than 50% (Supplementary Data Fig 3).

Fluorescence traces are presented in two normalized formats: ΔF/F and Norm F. ΔF/F is defined as:

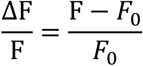

Where *F*_0_ is the average baseline fluorescence in the five seconds prior to stimulus onset. Norm F is defined as:

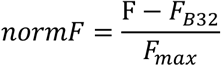

Where *F*_B32_ is the average baseline fluorescence in the five seconds prior to stimulus onset when the adaptation temperature is 32°C and *F_max_* is the maximum trial averaged fluorescence value across all stimulus conditions. Therefore, a Norm F value of 0 equates to a fluorescence value that matches the baseline fluorescence at an adaptation temperature of 32°C. A Norm F value of 1 equates to the maximum response from a given ROI.

#### Encoding bias parameter

The encoding bias parameter was estimated for each individual neuron based on the sensory evoked response across conditions. The goodness-of-fit to the absolute model (A) was estimated as the mean-squared-error relative to the mean response amplitude for each fixed TT across AT. The goodness-of-fit to the relative model (R) was estimated as the mean-squared-error relative to the mean response amplitude for each fixed ΔT across AT. The encoding bias parameter was defined as:

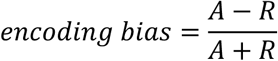

### Computational Model

#### Leaky integrate-and-fire model

A simple leaky integrate-and-fire (LIF) model was constructed with the addition of a cold-sensing (TRPM8) current. The LIF model is described by the following equations:

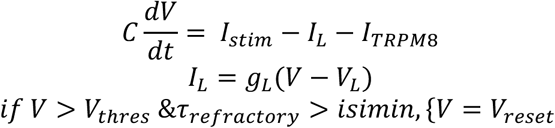

Model parameters were: V_L_ = -65 mV, C = 2µF/cm^2^, g_L_ = 0.035 mS/cm^2^, *V_thres_*=-35mV, *V_reset_* = -50 mV, *isimin*= 2 ms; as used previously^56–58^ ^3–5^. The TRPM8 current was defined as:

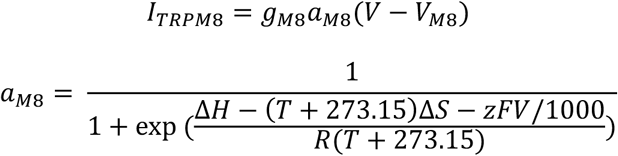

Model parameters were: Δ*H*=-156 kJ/mol, Δ*S*=-550J/molK, *z*=0.87, *R*=8.3144J/molK, *F*=96485C/mol; as used previously^41^. Thermosensory stimuli (T) were matched to experimental stimulus conditions to measure the simulated response. Model conductance (*g_MS_*) was varied within physiological ranges^42^ from 0.1-10 mS/cm^2^. Simulated neural responses were analyzed using the same techniques described for experimental data.

Experimental conductance values were estimated by goodness-of-fit for sensory evoked responses from the experimental neuron to the computationally modeled outputs across stimulus conditions. The *g*_/0_ value of the best-fit model, defined as the lowest means squared error, for each recorded neuron was defined as the estimated conductance.

## Supplementary Data

**Supplementary Data Figure 1.**
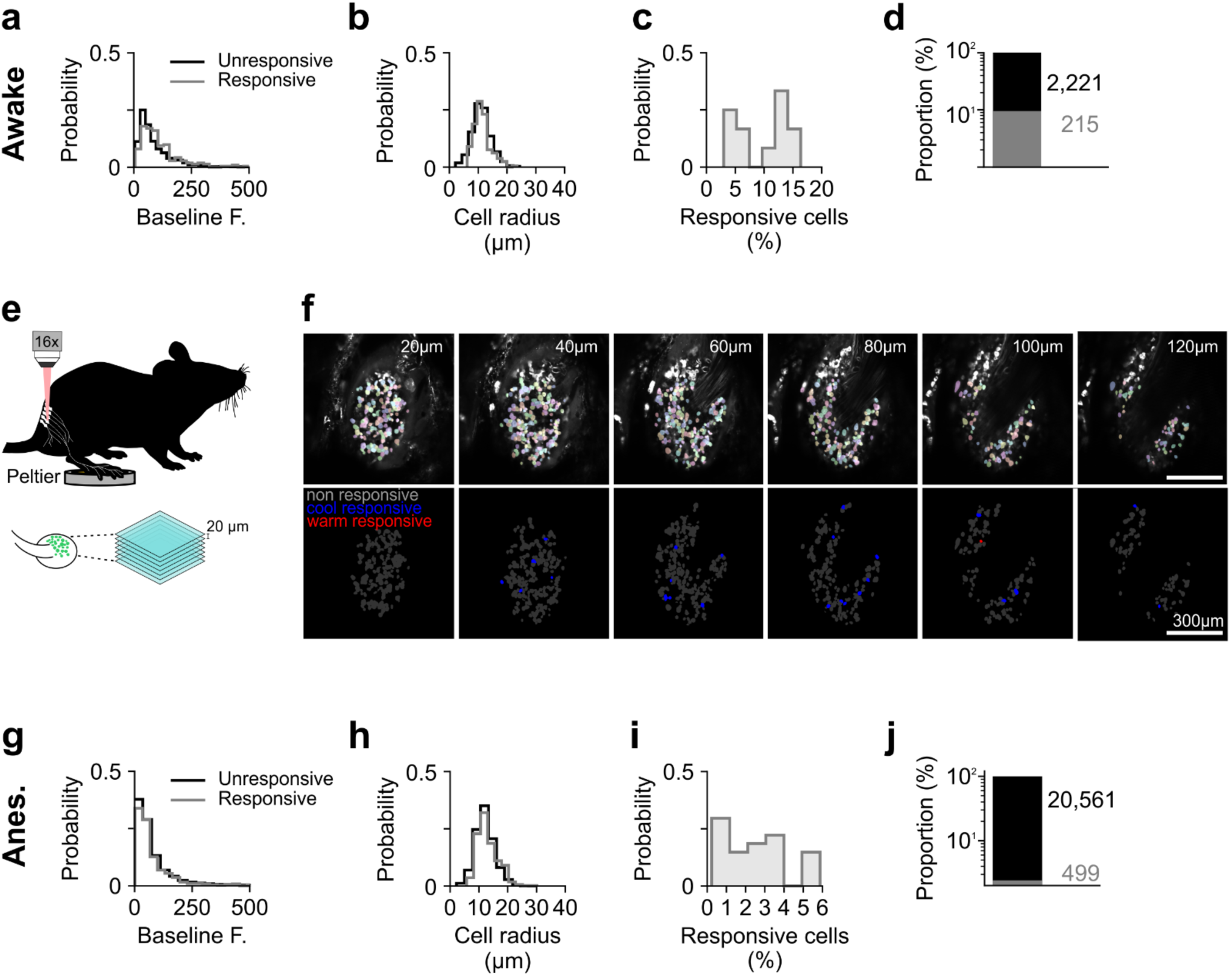
Thermal sensitivity is sparse across the dorsal root ganglion. **a.** Probability distribution of baseline fluorescence in awake mice for cells with (grey) or without (black) measured thermal sensitivity (n =2,436 neurons, 12 mice). **b.** Probability distribution of cell radius for cells in awake mice with (grey) or without (black) measured thermal sensitivity (*n =* 2,436 neurons, 12 mice). **c.** Probability distribution of percent of recorded ROIs in awake mice classified as responsive (*n =* 2,436 neurons, 12 mice). **d.** Proportion of anatomically identified ROIs in awake mice with (grey, *n =* 215 neurons) or without (black, *n =* 2,221 neurons) thermosensory evoked responses. **e.** Schematic depicting multi-plane imaging paradigm. **f**. Example of in vivo two-photon images from multi-plane sensory afferent neurons (top) with functional classification as cool responsive (blue) or warm responsive (red) overlaid on the image map (bottom). Scale bar, 300 µm. c. Probability distribution of baseline fluorescence for cells with (grey) or without (black) measured thermal sensitivity (*n =* 21,060 neurons, 30 mice). **g,h,i,j.** as a,b,c,d except under anesthesia (*n =* 21,060 total neurons, 20,561 unresponsive and 499 thermally responsive, 30 mice).

**Supplementary Data Figure 2.**
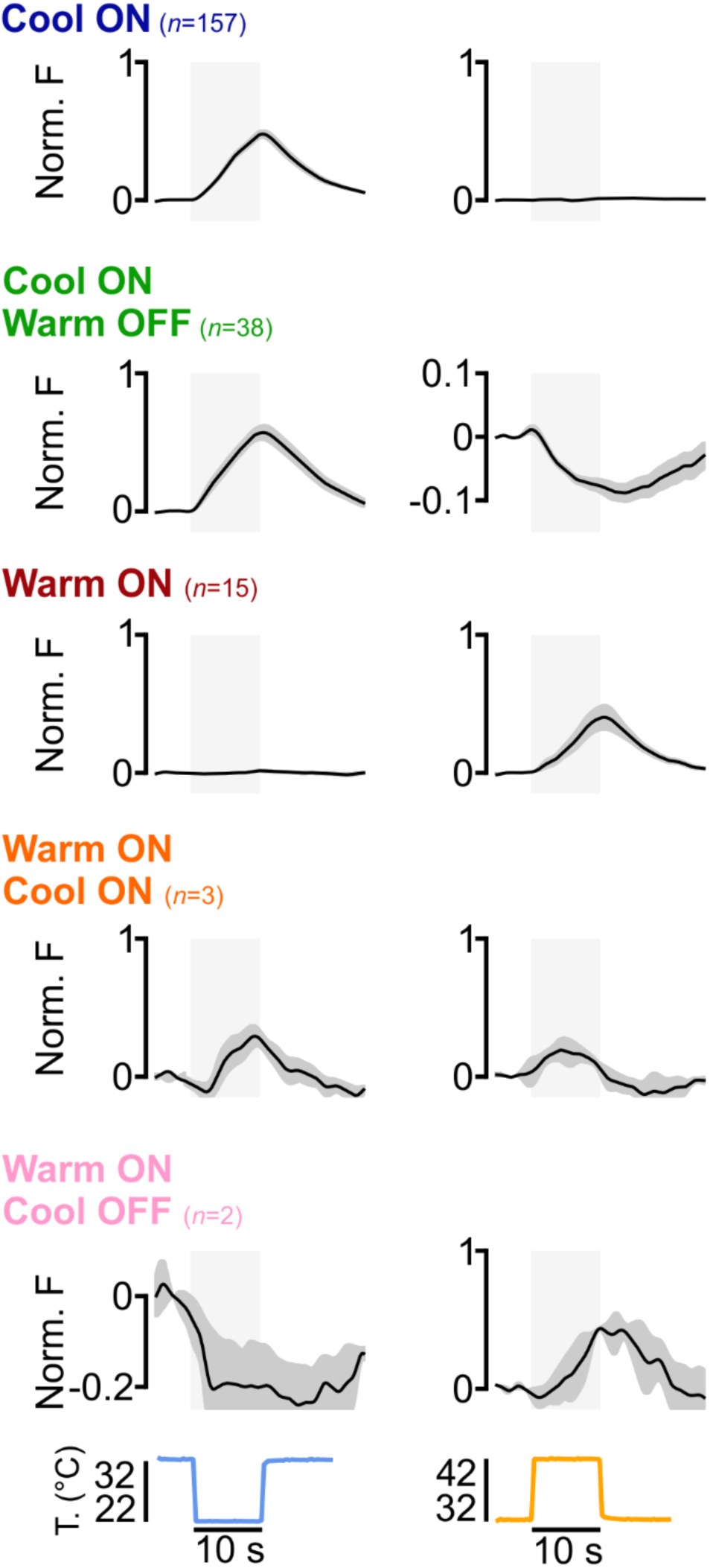
Thermosensory response profiles in thermoreceptors of the awake mouse. Grand average responses of thermoreceptor responses to cooling (left, 22°C) or warming (right, 42°C) thermal stimuli showing either an ON response to cooling (blue), an ON response to cooling and an OFF response to warming (green), an ON response to warming (maroon), an ON response to both warming and cooling (orange), or an ON response to warming and an OFF response to cooling (orange). Data shown as mean ± s.e.m..

**Supplementary Data Figure 3:**
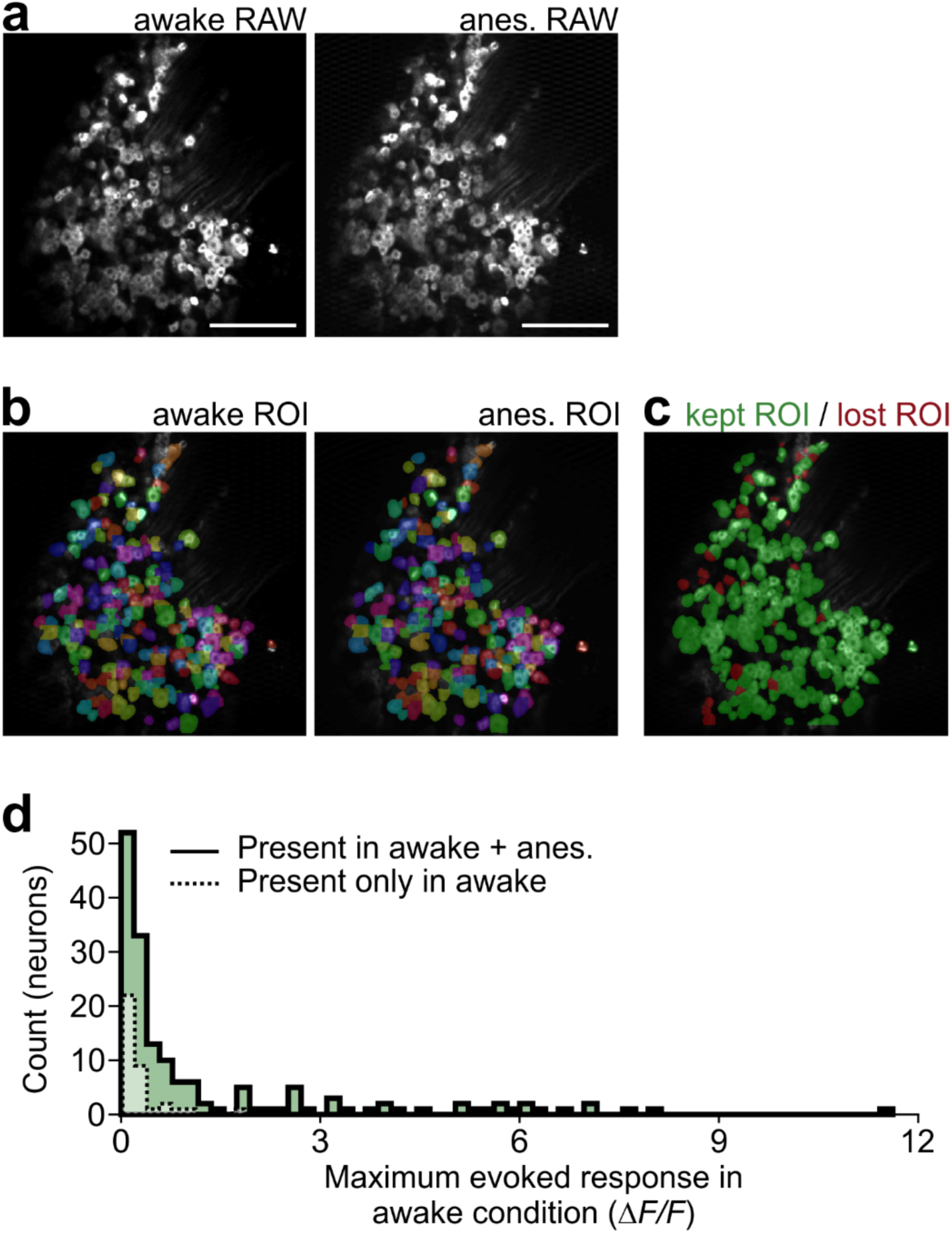
Methodological comparison of awake and anesthetized recordings. **a.** Average recorded image for the awake (left) and anesthetized (right) imaging session. **b.** Anatomical segmentation of the awake (left) and anesthetized (right) sessions using CellPose. Regions of interest (ROI) are pseudocolored and matched across each session. **c.** ROI segmentation of concatenated awake and anesthetized sesions using Suite2P. ROI are psuedocolored by those that are anatomically identified in both sessions and therefore kept for further analysis (green) and those that were not anatomically indentified in both sessions and therefore were excluded from further analysis (red). **d.** Neurons that were assessed as anatomically indentified in both sessions (green in panel c), but present only in the awake session (dotted outline) showed a lower stimulus evoked response in the awake condition than neurons that were functionally responsive in both sessions (solid outline).

**Supplementary Data Figure 4.**
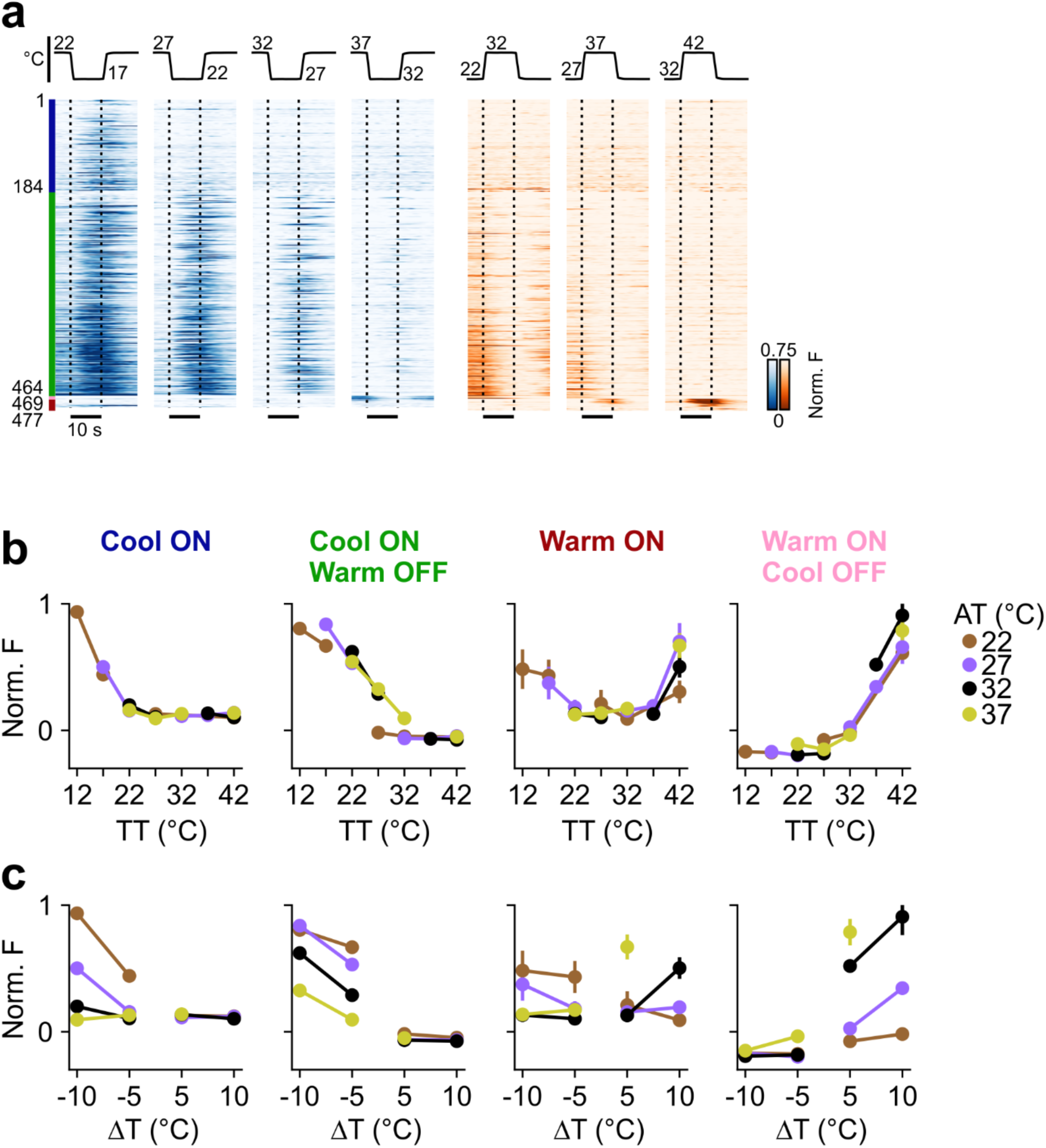
Absolute and relative encoding by thermoreceptors. **a.** Single-cell sensory afferent calcium responses to cooling (left, *ΔT* = -5°C) or warming (right, *ΔT* = +10°C) across AT. Each line represents a single neuron; sorted based on response latency and response type. Vertical dashed lines indicate the onset and end of the plateau phase of a thermal stimulus (*n =* 477 thermoreceptors, 16 mice). **b.** Normalized peak flourescence values by target temperature (TT) of each thermoreceptor identified. Color indicates the adaptation temperaure (AT = 22°C: brown, AT = 27°C: purple, AT = 32°C: black, AT = 37°C: yellow). **c.** Same as b, but depicted as a change in temperature (ΔT). Data shown as mean ± s.e.m..

**Supplementary Data Figure 5.**
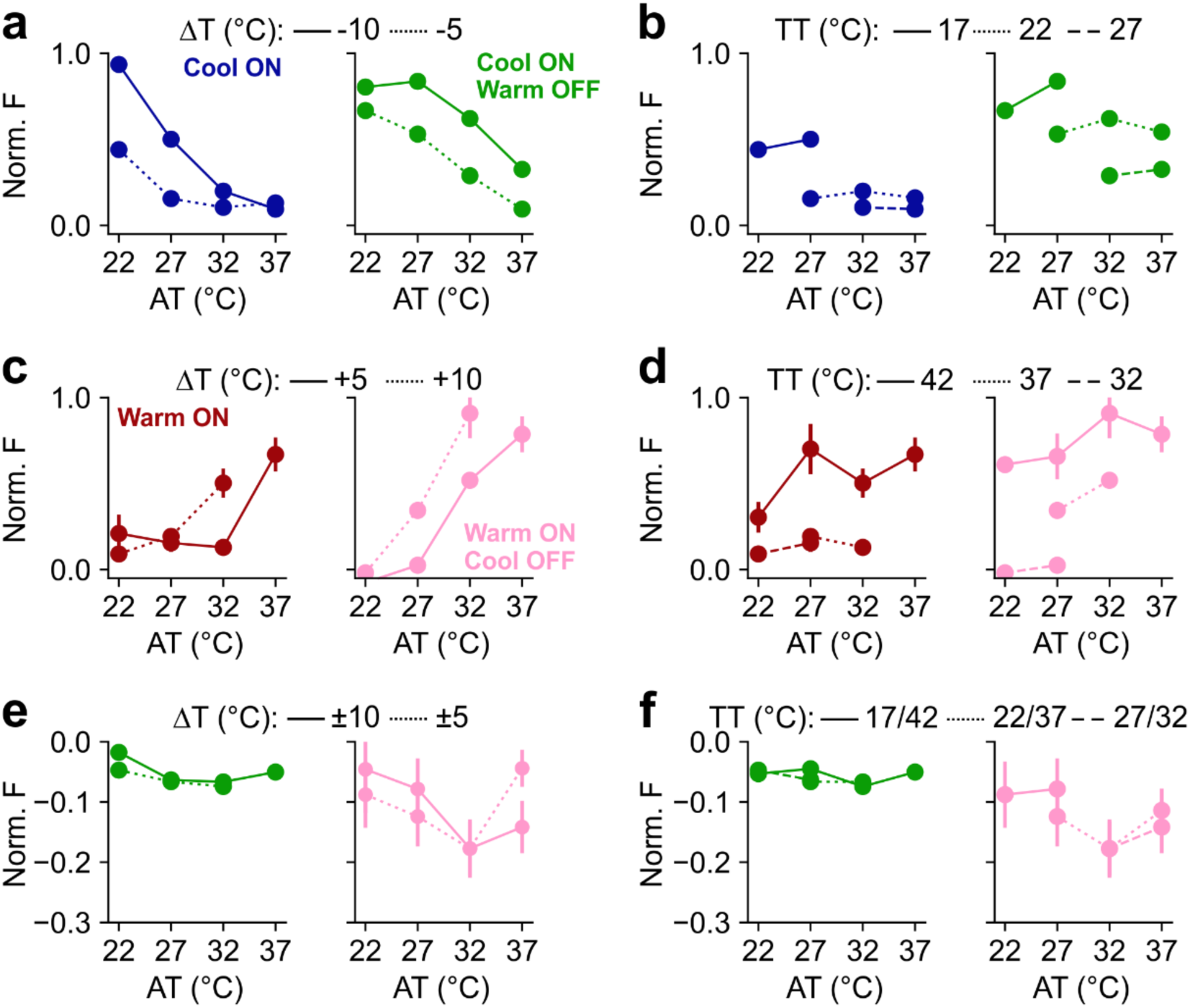
ON/OFF cells show greater sensitivity than ON-only cells. **a.** Normalised peak fluorescence response values across AT in response to a ΔT of -10°C (solid) or -5°C (dotted) for Cool-ON (left: *n =* 184 cells, 16 mice, blue) and Cool-ON/Warm-OFF (right: *n =* 280 cells, 16 mice, green). **b.** Same as a except for TT = 17°C (solid), 22°C (dotted), or 27°C (dashed). **c.** Same as a except for ΔT = +5°C (solid) or +10°C (dotted) for Warm-ON (left: *n =* 8 cells, 16 mice, maroon) and Warm-ON/Cool-OFF (right: *n =* 5 cells, 16 mice, pink). **d.** Same as b except for TT = 42°C (solid), 37°C (dotted), or 32°C (dashed). **e.** Same as a/c, but for the OFF responses for warming (left: Cool-ON/Warm-OFF, green, ΔT = +5°C (dotted) or +10°C (solid)) and cooling (right: Warm-ON/Cool-OFF, pink, ΔT = -5°C (dotted) or -10°C (solid)). **f.** Same as for e, but for TT = 17/42°C (solid), 22/37°C (dotted), or 27/32°C (dashed). Data shown as mean ± s.e.m..

**Supplementary Data Figure 6.**
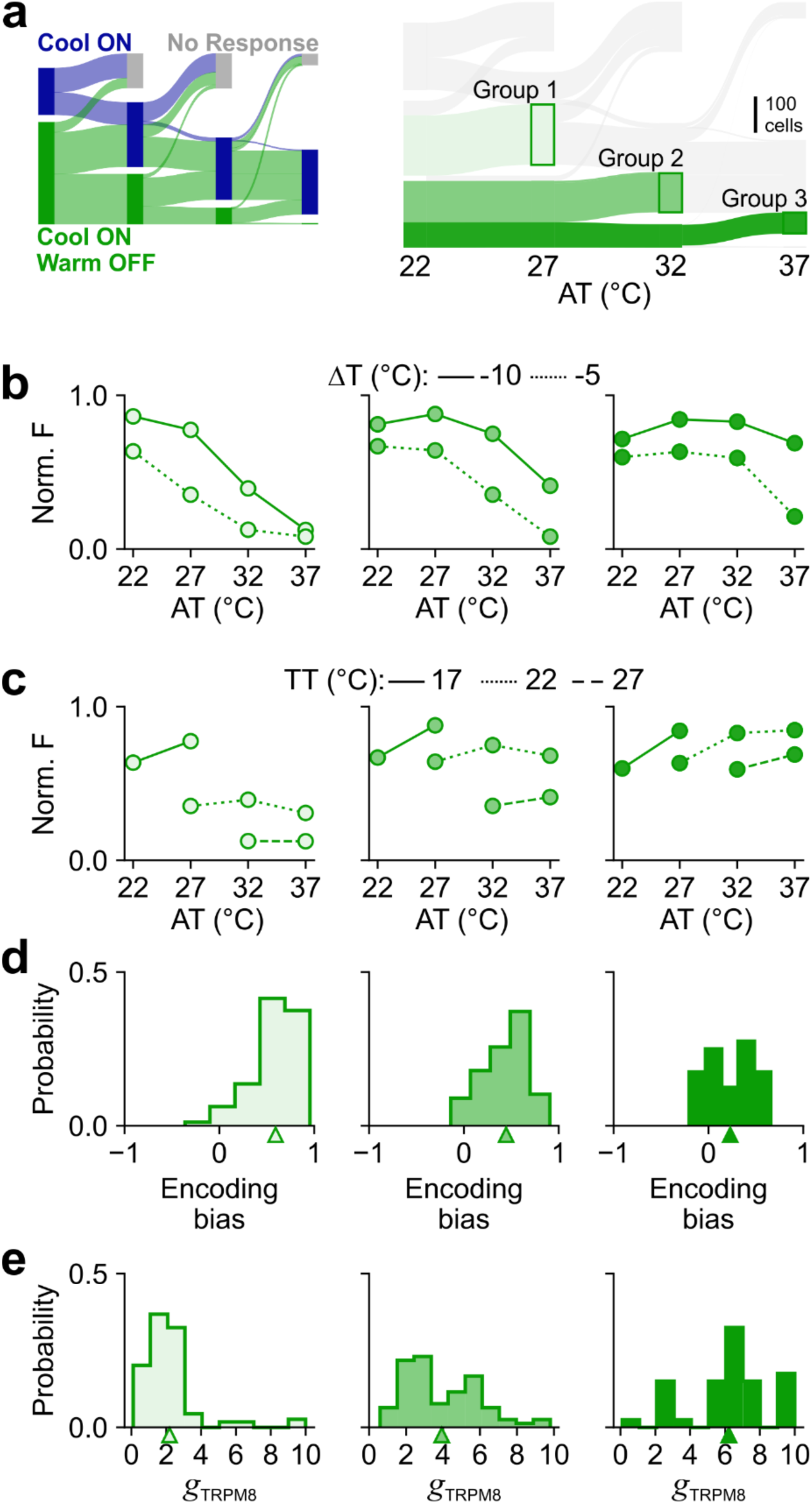
Retention of a Warm OFF response at higher AT indicates higher sensitivity. **a.** Left. Sankey plot depicting transitions of neurons between functionally defined Cool-ON thermosensory response profiles over all AT. Responsive cells are classified according to thermal tuning preference, with neurons classified as none if no temperature evoked calcium transients were observed. Bar colour indicates classification at each AT while ribbon colour indicates classification at the lowest AT (22°C). Right. Sankey plot of only the Cool-ON/Warm-OFF responses with group identification indicating the AT at which the Warm OFF response was lost. Group 1 Cool-ON/Warm-OFF cells no longer displayed an OFF response at 27°C AT. Group 2 maintained an OFF response at 27°C AT but did not at 32°C AT. Group 3 maintained the warm OFF response at 32°C AT. **b.** Normalised peak fluorescence response values across AT in response to a ΔT of -10°C (solid) or -5°C (dotted) for Group 1-3 (left to right). **c.** Same as c, except for TT = 17°C (solid), 22°C (dotted), or 27°C (dashed). **d.** Encoding bias, defined as the similarity of an individual unit tuning to absolute (+1) or relative (-1) encoding in response to cooling for Group 1-3 (left to right). **e.** Model-fit TRPM8 conductance values to each experimental neuron in Group 1 (left: *n =* 114 neurons), Group 2 (middle: *n =* 78 neurons), and Group 3 (right: *n =* 40 neurons). Data shown as mean ± s.e.m..

## Notes

### Competing Interest Statement

The authors have declared no competing interest.

## References

1. Zotterman, Y. Specific action potentials in the lingual nerve of cat. Skand. Arch. Für Physiol. 75, 105–119 (1936).

2. Blix, M. Experimentela bidrag till losning af fragan om hudnervernas specifika energi. Upps. Lakfor Forh 87–102 (1882).

3. Vriens, J., Nilius, B. & Voets, T. Peripheral thermosensation in mammals. Nat. Rev. Neurosci. (2014) doi:10.1038/nrn3784.

4. Filingeri, D. Neurophysiology of Skin Thermal Sensations. in Comprehensive Physiology (ed. Prakash, Y. S.) 1429–1491 (Wiley, 2016). doi:10.1002/cphy.c150040.

5. Stevens, J. C. & Choo, C. Temperature sensitivity of the body surface over the life span. Somatosens. Mot. Res. 15, 13–28 (1998).

6. Frenzel, H. et al. A Genetic Basis for Mechanosensory Traits in Humans. PLoS Biol. 10, e1001318 (2012).

7. Paricio-Montesinos, R. et al. The Sensory Coding of Warm Perception. Neuron 106, 830–841.e3 (2020).

8. Darian-Smith, I., Johnson, K. O. & Dykes, R. ‘Cold’ fiber population innervating palmar and digital skin of the monkey: responses to cooling pulses. J. Neurophysiol. 36, 325–346 (1973).

9. LaMotte, R. H. & Campbell, J. N. Comparison of responses of warm and nociceptive C-fiber afferents in monkey with human judgments of thermal pain. J. Neurophysiol. 41, 509– 528 (1978).

10. Darian-Smith, I. et al. Warm fibers innervating palmar and digital skin of the monkey: responses to thermal stimuli. J. Neurophysiol. 42, 1297–1315 (1979).

11. Hallin, R. G., Torebjork, H. E. & Wiesenfeld, Z. Nociceptors and warm receptors innervated by C fibres in human skin. J. Neurol. Neurosurg. Psychiatry 45, 313–319 (1982).

12. Milenkovic, N. et al. A somatosensory circuit for cooling perception in mice. Nat. Neurosci. 17, 1560–1566 (2014).

13. Bokiniec, P., Zampieri, N., Lewin, G. R. & Poulet, J. F. The neural circuits of thermal perception. Curr. Opin. Neurobiol. 52, 98–106 (2018).

14. Vestergaard, M., Carta, M., Güney, G. & Poulet, J. F. A. The cellular coding of temperature in the mammalian cortex. Nature 614, 725–731 (2023).

15. Leva, T., et al. The spatial representation of temperature in the thalamus. bioRxiv (2024).

16. Hensel, H. & Iggo, A. Analysis of cutaneous warm and cold fibres in primates. Pfluegers Arch. Eur. J. Physiol. 329, 1–8 (1971).

17. Yarmolinsky, D. A. et al. Coding and Plasticity in the Mammalian Thermosensory System. Neuron 92, 1079–1092 (2016).

18. McKemy, D. D., Neuhausser, W. M. & Julius, D. Identification of a cold receptor reveals a general role for TRP channels in thermosensation. Nature 416, 52–58 (2002).

19. Knowlton, W. M. et al. A sensory-labeled line for cold: TRPM8-expressing sensory neurons define the cellular basis for cold, cold pain, and cooling-mediated analgesia. J. Neurosci. (2013) doi:10.1523/jneurosci.1943-12.2013.

20. Voets, T. TRP Channels and Thermosensation. in Mammalian Transient Receptor Potential (TRP) Cation Channels (eds. Nilius, B. & Flockerzi, V.) vol. 223 729–741 (Springer International Publishing, Cham, 2014).

21. Palkar, R., Lippoldt, E. K. & McKemy, D. D. The molecular and cellular basis of thermosensation in mammals. Curr. Opin. Neurobiol. 34, 14–19 (2015).

22. Pogorzala, L. A., Mishra, S. K. & Hoon, M. A. The Cellular Code for Mammalian Thermosensation. J. Neurosci. 33, 5533–5541 (2013).

23. Hensel, H., Iggo, A. & Witt, I. A quantitative study of sensitive cutaneous thermoreceptors with C afferent fibres. J. Physiol. 153, 113–126 (1960).

24. Kenshalo, D. R. & Gallegos, E. S. Multiple Temperature-Sensitive Spots Innervated by Single Nerve Fibers. Science 158, 1064–1065 (1967).

25. Dubner, R., Sumino, R. & Wood, W. I. A peripheral ‘cold’ fiber population responsive to innocuous and noxious thermal stimuli applied to monkey’s face. J. Neurophysiol. 38, 1373–1389 (1975).

26. Gallio, M., Ofstad, T. A., Macpherson, L. J., Wang, J. W. & Zuker, C. S. The Coding of Temperature in the Drosophila Brain. Cell 144, 614–624 (2011).

27. Campero, M., Serra, J., Bostock, H. & Ochoa, J. L. Slowly conducting afferents activated by innocuous low temperature in human skin. J. Physiol. 535, 855–865 (2001).

28. LaMotte, R. H. & Thalhammer, J. G. Response properties of high-threshold cutaneous cold receptors in the primate. Brain Res. 244, 279–287 (1982).

29. Wang, F. et al. Sensory Afferents Use Different Coding Strategies for Heat and Cold. Cell Rep. 23, 2001–2013 (2018).

30. Emery, E. C. et al. In vivo characterization of distinct modality-specific subsets of somatosensory neurons using GCaMP. Sci. Adv. 2, e1600990 (2016).

31. Chisholm, K. I., Khovanov, N., Lopes, D. M., La Russa, F. & McMahon, S. B. Large Scale *In Vivo* Recording of Sensory Neuron Activity with GCaMP6. eneuro 5, ENEURO.0417–17.2018 (2018).

32. Qi, L. et al. A mouse DRG genetic toolkit reveals morphological and physiological diversity of somatosensory neuron subtypes. Cell 187, 1508–1526.e16 (2024).

33. Mishra, S. K., Tisel, S. M., Orestes, P., Bhangoo, S. K. & Hoon, M. A. TRPV1-lineage neurons are required for thermal sensation. EMBO J. 30, 582–593 (2011).

34. Chen, C. et al. Long-term imaging of dorsal root ganglia in awake behaving mice. Nat. Commun. 10, 3087 (2019).

35. Ran, C., Hoon, M. A. & Chen, X. The coding of cutaneous temperature in the spinal cord. Nat. Neurosci. 19, 1201–1209 (2016).

36. MacIver, M. B. & Tanelian, D. L. Volatile anesthetics excite mammalian nociceptor afferents recorded in vivo. Anesthesiology (1990).

37. Raithel, S. J., Sapio, M. R., Iadarola, M. J. & Mannes, A. J. Thermal A-δ Nociceptors, Identified by Transcriptomics, Express Higher Levels of Anesthesia-Sensitive Receptors Than Thermal C-Fibers and Are More Suppressible by Low-Dose Isoflurane. Anesth. Analg. 127, 263–266 (2018).

38. Zaproudina, N., Varmavuo, V., Airaksinen, O. & Närhi, M. Reproducibility of infrared thermography measurements in healthy individuals. Physiol. Meas. 29, 515–524 (2008).

39. Xiao, R., Xiao, R., Xu, X. Z. S. & Xu, X. Z. S. Temperature sensation from molecular thermosensors to neural circuits and coding principles. Annu. Rev. Physiol. (2021) doi:10.1146/annurev-physiol-031220-095215.

40. Knowlton, W. M., Daniels, R. L., Palkar, R., McCoy, D. D. & McKemy, D. D. Pharmacological Blockade of TRPM8 Ion Channels Alters Cold and Cold Pain Responses in Mice. PLoS ONE 6, e25894 (2011).

41. McGahan, K. & Keener, J. A mathematical model analyzing temperature threshold dependence in cold sensitive neurons. PLOS ONE 15, e0237347 (2020).

42. Olivares, E. et al. TRPM8-Dependent Dynamic Response in a Mathematical Model of Cold Thermoreceptor. PLOS ONE 10, e0139314 (2015).

43. Voets, T. et al. The principle of temperature-dependent gating in cold- and heat-sensitive TRP channels. Nature 430, 748–754 (2004).

44. Madrid, R., De La Peña, E., Donovan-Rodriguez, T., Belmonte, C. & Viana, F. Variable Threshold of Trigeminal Cold-Thermosensitive Neurons Is Determined by a Balance between TRPM8 and Kv1 Potassium Channels. J. Neurosci. 29, 3120–3131 (2009).

45. Lemon, C. H., Kang, Y. & Li, J. Separate functions for responses to oral temperature in thermo-gustatory and trigeminal neurons. Chem. Senses 41, 457–471 (2016).

46. Li, J., Zumpano, K. T. & Lemon, C. H. Separation of Oral Cooling and Warming Requires TRPM8. J. Neurosci. 44, e1383232024 (2024).

47. Iggo, A. Cutaneous thermoreceptors in primates and sub-primates. J. Physiol. 200, 403– 430 (1969).

48. El Hay, M. Y. A., Kamm, G. B., Tlaie, A. & Siemens, J. Diverging roles of TRPV1 and TRPM2 in warm-temperature detection. Preprint at 10.7554/eLife.95618.1 (2024).

49. Vilar, B., Tan, C.-H. & McNaughton, P. A. Heat detection by the TRPM2 ion channel. Nature 584, E5–E12 (2020).

50. Chisholm, K. I. et al. Encoding of cutaneous stimuli by lamina I projection neurons. Pain 162, 2405–2417 (2021).

51. Liu, W. W., Mazor, O. & Wilson, R. I. Thermosensory processing in the Drosophila brain.Nature 519, 353–357 (2015).

52. Alpert, M. H. et al. A Circuit Encoding Absolute Cold Temperature in Drosophila. Curr. Biol. 30, 2275–2288.e5 (2020).

53. Daigle, T. L. et al. A Suite of Transgenic Driver and Reporter Mouse Lines with Enhanced Brain-Cell-Type Targeting and Functionality. Cell 174, 465–480.e22 (2018).

54. Stringer, C., Wang, T., Michaelos, M. & Pachitariu, M. Cellpose: a generalist algorithm for cellular segmentation. Nat. Methods 18, 100–106 (2021).

55. Pachitariu, M. et al. Suite2p: beyond 10,000 neurons with standard two-photon microscopy. Preprint at 10.1101/061507 (2016).

56. Whitmire, C. J., Waiblinger, C., Schwarz, C. & Stanley, G. B. Information Coding through Adaptive Gating of Synchronized Thalamic Bursting. Cell Rep. 14, 795–807 (2016).

57. Ladenbauer, J., Augustin, M., Shiau, L. & Obermayer, K. Impact of Adaptation Currents on Synchronization of Coupled Exponential Integrate-and-Fire Neurons. PLoS Comput. Biol. 8, e1002478 (2012).

58. Lesica, N. A. Encoding of Natural Scene Movies by Tonic and Burst Spikes in the Lateral Geniculate Nucleus. J. Neurosci. 24, 10731–10740 (2004).

